# Circular RNA aptamers as superb sub-stoichiometric chaperones of TDP-43 condensation

**DOI:** 10.64898/2026.06.01.729436

**Authors:** Hao Wu, Peng-Fei Luan, Jiaojiao Hu, Fang Nan, Ling Li, Yu-Yin Pan, Bo-Wen Jiang, Jie-Wen Chen, Yu-Xin Liu, Yichen Wei, Xiao Wang, Jiaquan Liu, Guang Xu, Cong Liu, Ling-Ling Chen

**Affiliations:** State Key Laboratory of RNA Innovation, Science and Engineering, New Cornerstone Science Laboratory, CAS Center for Excellence in Molecular Cell Science, Shanghai Institute of Biochemistry and Cell Biology, University of Chinese Academy of Sciences, Chinese Academy of Sciences, Shanghai 200031, China; Interdisciplinary Research Center on Biology and Chemistry, State Key Laboratory of Chemical Biology, Shanghai Institute of Organic Chemistry, Chinese Academy of Sciences, Shanghai 201210, China; School of Life Science and Technology, ShanghaiTech University, Shanghai 201210, China; School of Life Science and Biotechnology, Shanghai Jiao Tong University, Shanghai 200240, China; Shanghai Academy of Natural Sciences (SANS), Shanghai 200031, China

## Abstract

Pathological protein misfolding and aggregation underlie numerous human diseases, yet strategies that selectively antagonize pathological protein states while preserving normal protein function remain limited. Here, we identify two loosely structured circular RNA aptamers (Capts), circT3 and its minimized 116-nt derivative circT3-M3, as superb sub-stoichiometric RNA chaperones that suppress pathological condensation of TAR DNA-binding protein 43 (TDP-43), a hallmark of amyotrophic lateral sclerosis and frontotemporal dementia. Rather than acting through simple stoichiometric sequestration, these Capts engage TDP-43 reversibly and iteratively via binding the functionally and pathologically relevant residues within its RRM1 domain. Such interaction enables a single Capt to promote rapid and multiple RRM1-dependent homomeric assembly events, thereby stabilizing TDP-43 in a soluble oligomeric state to outcompete the C-terminal prion-like domain-driven condensation and supersaturation. Moreover, delivery of these Capts rescues TDP-43-dependent aberrant splicing in the cellular model and partially ameliorates TDP-43 proteinopathy-driven motor dysfunction *in vivo*. Together, our findings establish Capts as highly efficient subcellular RNA chaperones with exceptionally potent low-dose activity and highlight their promise as next-generation RNA therapeutics for TDP-43 proteinopathies.

## INTRODUCTION

Protein conformational conversion, rather than altered expression alone, is increasingly recognized as a central driver of many diseases^1,2^. In the nervous system, prion-like domain (PrLD)-containing proteins, including TAR DNA-binding protein 43 (TDP-43) and FUS, are particularly prone to liquid-liquid phase separation (LLPS) under low-RNA conditions^3^. During pathological stress, these proteins mislocalize from the RNA-rich nucleus to the relatively RNA-poor cytoplasm, where they undergo LLPS, form condensates, and eventually mature into solid-like aggregates^3^. Pathological TDP-43 aggregation is a defining feature of amyotrophic lateral sclerosis (ALS) and frontotemporal dementia (FTD), occurring in approximately 97% of ALS cases and nearly half of FTD cases^4,5^. Mechanistically, TDP-43 supersaturation combined with oxidative stress drives demixing within condensates, yielding a TDP-43-enriched phase that progressively solidifies into pathological aggregates^6^. Thus, preventing TDP-43 aggregation is considered as a promising therapeutic strategy to slow or ameliorate the progression of TDP-43-associated neurodegenerative diseases^7,8^. However, selectively eliminating pathological TDP-43 assemblies at the subcellular level while preserving the physiological pool of TDP-43 remains a major challenge.

In addition to its C-terminal PrLD, also known as the C-terminal low-complexity domain (LCD), TDP-43 contains a globular N-terminal domain (NTD) and two RNA recognition motifs (RRM1 and RRM2). Under physiological conditions, nuclear TDP-43 forms homomeric oligomers via its NTD^9^ and binds to 3′ untranslated regions (3′ UTRs) or intronic sequences of target RNAs to regulate alternative splicing and gene expression^10,11^. These bound RNAs in turn serve as scaffolds that promote TDP-43 oligomerization^3^. Critically, this oligomerization spatially segregates the adjacent and highly aggregation-prone C-terminal domain (CTD), thereby restraining its tendency to form pathological aggregates^9^. Together, promoting or stabilizing physiological TDP-43 oligomerization emerges as a promising strategy to counteract pathological condensation and aggregation.

RNA aptamers offer a versatile platform for modulating pathological proteins that are difficult to target with conventional small molecule-or protein-based therapeutics^12^. However, their therapeutic potential is limited by challenges such as susceptibility to RNase degradation and immunogenicity. A recent study showed that short RNAs, including *Clip34* and *Malat1_start* derived from the lncRNA *Malat1*, promote aggregation-resistant TDP-43 conformers by allosterically destabilizing a conserved helical region in its prion-like domain, suggesting RNA aptamers could prevent pathological TDP-43 aggregation^13^. However, this model requires RNA molecules exceeding TDP-43, limiting its medical applicability^13^. Circular RNAs, which exhibit high stability^14,15^, distinctive conformational properties^16,17^, and low immunogenicity^18,19^, have emerged as promising alternatives to linear aptamers (Lapts). Our previous studies have shown that circular RNA aptamers (Capts) containing 16-26 bp imperfect duplexes (ds-cRNAs) can suppress aberrant activation of double-stranded RNA-activated protein kinase R (PKR)^16,18^ and ameliorate pathological phenotypes in mouse models of psoriasis, osteoarthritis, and Alzheimer’s disease^19–21^. This subcellular precise targeting of pathological PKR activation by Capts promoted us to hypothesize that Capts may also be applicable to specifically eliminate proteins involved in other pathological contexts, such as pathological TDP-43 aggregates^22^.

Although Capts possess distinct advantages, minimizing the required dosage remains important for reducing potential side effects. Thus, when designing aptamers to target pathological TDP-43, identifying candidates effective at low concentrations is necessary. Intriguingly, the long noncoding RNA *SLERT* was found to act as an RNA chaperone to modulate the RNA helicase DDX21 conformation at a remarkably low stoichiometric ratio (∼ 1 RNA molecule per 1,000 protein molecules) within the nucleolus^23^. This observation prompted us to ask the possibility to develop specific Capts that could specifically prevent pathological TDP-43 aggregation at similarly low stoichiometric ratios, as a new therapeutic strategy to ameliorate the progression of TDP-43-associated neurodegenerative diseases (Fig. 1a).

**Fig. 1.**
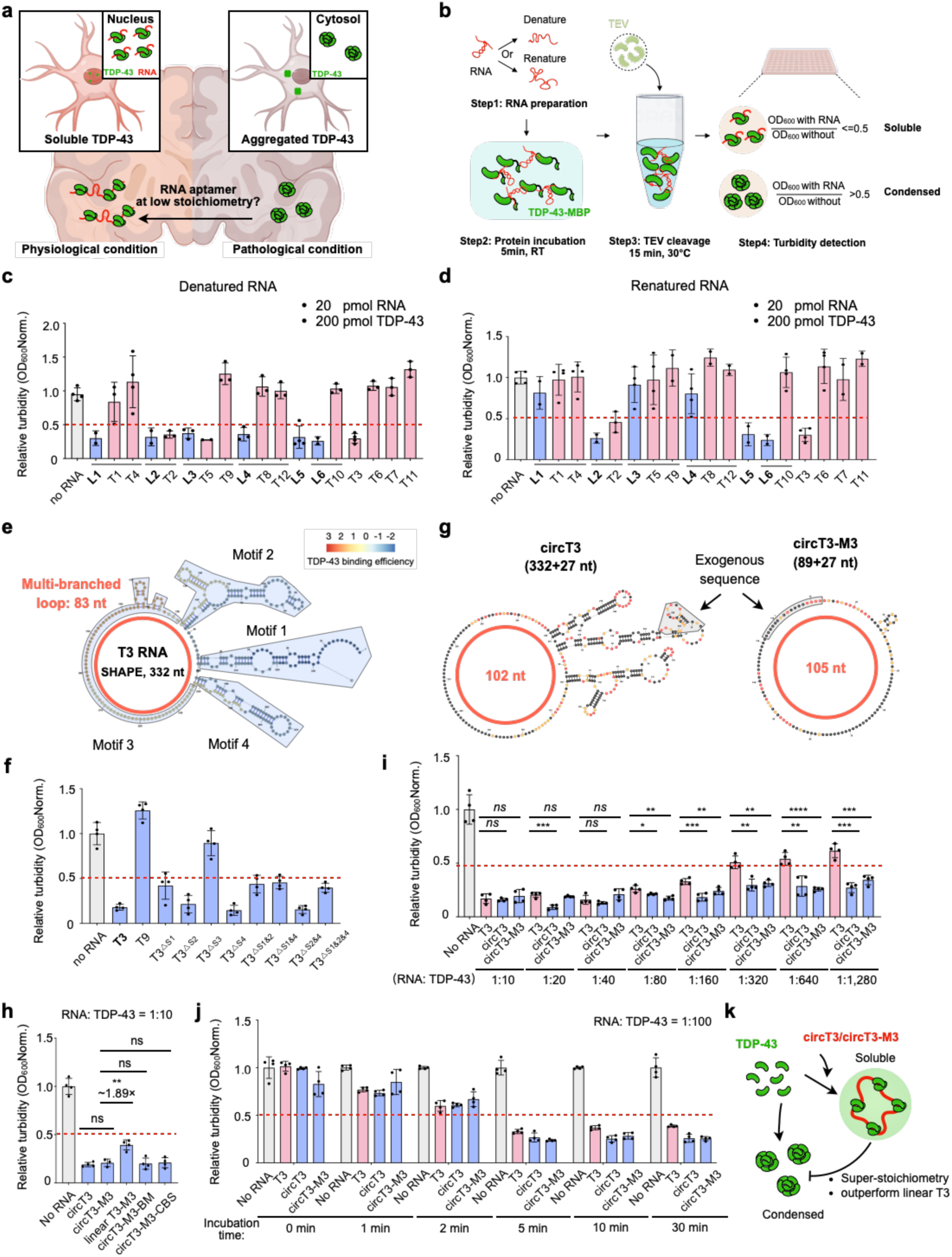
Circular RNA aptamers suppress TDP-43 condensation at superb sub-stoichiometric ratios. **a** Conceptual schematic of this study. In physiological condition, TDP-43 localizes to the nucleus of neuronal cells and remains in soluble states. Upon pathological stress, TDP-43 is exported into the cytoplasm and forms aggregation. In this study, we aim to identify RNA aptamers that suppress TDP-43 pathological aggregation at low stoichiometry. **b** Schematic of the turbidity assay used to detect whether the RNA fragments can suppress TDP-43 condensation *in vitro*. Both the denatured and renatured RNAs are applied for evaluation. The RNAs are pre-incubated with C-terminal MBP-tagged TDP-43 oligomer that harbors an internal TEV cleavage site. After TEV treatment to remove the MBP tag, the Turbidity of the samples are detected to evaluate the condensation potential of TDP-43. **c** Turbidity assay shows that denatured L1-L6, T2, T3, and T5 RNA fragments significantly reduced TDP-43 condensation *in vitro*. **d** Turbidity assay shows that renatured L2, L5, L6, T2, and T3 RNA fragments suppressed TDP-43 condensation *in vitro*. **e** Secondary structure of T3 RNA modeled with SHAPE-MaP experiment. Three duplex-containing structures (Motif 1, 2, 4) and the multi-branched loop structure (Motif 3, 83-nt in length) are marked in blue. TDP-43 binding efficiency is indicated from the TDP-43 iCLIP-seq data. **f** Deletion the Motif 3 but not the duplex-containing structures completely abolished the efficacy of T3 in suppressing TDP-43 condensation, while deletion of Motif 1 has mild effect. T9 RNA is used as negative control. **g** Secondary structures of circT3 and circT3-M3 modeled with SHAPE-MaP experiments. circT3 is 359-nt in length containing a 102-nt multi-branched loop structure, circT3-M3 is 116-nt in length containing a 105-nt multi-branched loop structure. The exogenous 27 nt sequence generated during circularizing process is marked in grey. **h** circT3-M3 shows comparable efficacy to circT3 and ∼1.89-fold efficacy to linear T3-M3 in suppressing TDP-43 condensation. Addition of the TDP-43 binding sequence (BM) or core binding sequence (CBS) does not improve this efficacy, revealed by the turbidity assay. Oligomeric MBP-tagged TDP-43 and renatured RNAs are used. **i** The RNA aptamers show efficacy in suppressing TDP-43 condensation sub-stoichiometrically. Oligomeric MBP-tagged TDP-43 and renatured RNAs are used in this experiment. The RNA: TDP-43 molar ratios varying from 1:10 to 1:1,280 are evaluated. Capts outperform linear T3 at low stoichiometry, revealed by the turbidity assay. **j** At an RNA: TDP-43 molar ratio of 1:100, the RNA aptames begin to exhibit detectable efficacy in preventing TDP-43 condensation at 5 min of co-incubation. In this study, the renatured RNA aptamers are pre-incubated with MBP-tagged TDP-43 oligomers for 0, 1, 2, 5, 10, 20 min. Then the mixed samples are subject to 5 min TEV enzyme treatment followed by turbidity detection. **k** Schematic shows that Capts with large and loosened structures prevent TDP-43 condensation superb sub-stoichiometrically and outperforms their linear counterparts.

Here, we developed Capts, circT3 and its minimized 116-nt derivative circT3-M3, to efficiently suppress TDP-43 condensation, an intermediate state that underlies most pathological aggregation^6^. These Capts acted as RNA chaperones that transiently and iteratively engaged functionally and pathologically relevant residues within the TDP-43 RRM1 domain at an ultralow stoichiometric ratio. Such interaction promoted RRM1-dependent TDP-43 homomeric assembly and favored soluble TDP-43 oligomers over C-terminal PrLD-driven condensation and supersaturation. Importantly, the LNP-delivered Capts reduced cytosolic TDP-43 condensation and restored TDP-43-dependent aberrant splicing in cellular models. AAV-mediated expression of either Capt prevented cytosolic TDP-43 accumulation and partially ameliorated motor dysfunction in the TDP-43 proteinopathy mouse model. Together, these findings reveal that Capts can act as robust subcellular modulators of pathological TDP-43 conformations and suggest a potential RNA-based therapeutic strategy to resolve TDP-43-associated proteinopathies.

## RESULTS

### RNAs with loosened structures inhibit TDP-43 condensation *in vitro*

To identify RNAs capable of preventing pathological TDP-43 assembly, we reasoned that candidate RNAs should bind TDP-43. Analysis of TDP-43 iCLIP-seq datasets from human embryonic stem cells (H9) (GSM2285561 and GSM2285562)^24^ identified 1,646 binding sites at a false discovery rate (FDR) below 0.05. These sites were clustered into RNA fragments using distance cutoffs of ≤100 nt or ≤200 nt, yielding 255 and 236 fragments (designated Candidate Region 1 and 2; Supplementary Fig. S1a), respectively. From these pools, we selected the top 19 fragments, including 12 from Candidate Region 1 (T1-T12) and 7 from Candidate Region 2 (L1-L7) for further analyses (Supplementary Fig. S1a). These selected fragments ranged from 136 to 959 nt, several contained known TDP-43-binding motifs, such as UG repeats^25^, but no other clear sequence preference was apparent. Most of them were originated from the lncRNAs *NEAT1* and *MALAT1* or from mitochondrial RNAs (Supplementary Fig. S1b, c), consistent with reported roles of TDP-43 in RNA splicing^10,26^ and respiratory complex I assembly^27^.

Because TDP-43 preferentially binds single-stranded RNAs (ssRNAs)^28^, we hypothesized that structured RNAs would bind TDP-43 less efficiently and would therefore be less effective at suppressing TDP-43 condensation. To test this, we prepared each RNA under denaturing (less structured) or renaturing (more structured) conditions (Fig. 1b) and incubated it with C-terminal MBP-tagged TDP-43 oligomers containing an internal TEV-cleavage site at an RNA: TDP-43 molar ratio of 1:10. After TEV cleavage to remove the MBP tag, the resulting mixtures were analyzed by turbidity assay to dissect their condensation potential (Fig. 1b). Under denaturing conditions, we found that the RNA fragments L1-L6, T2, T3, and T5 remarkably reduced TDP-43 condensation (Fig. 1c). However, after renaturation in 10 mM magnesium, which promoted RNA folding, L1, L3, L4, and T5 RNAs lost detectable inhibitory activity (Fig. 1d). In addition, removing magnesium or replacing it with potassium, but not calcium, restored the activity of L4 and T5 (Supplementary Fig. S1d), indicating that RNA folding state directly influenced anti-condensation activity. These results suggest that TDP-43-interacting RNAs with minimal secondary structures are more effective at inhibiting TDP-43 condensation *in vitro*.

Given that intracellular free magnesium concentration is estimated at ∼1 mM^29^, we asked whether such Mg²⁺ levels would affect RNA activity. Although the L4 activity was reduced at 10 mM Mg²⁺, it retained robust anti-TDP-43 condensation efficacy at 1 mM Mg²⁺, comparable to that of L6 and T3 (Supplementary Fig. S1e). These observations indicate that the tested RNA fragments can remain functional in suppressing TDP-43 condensation under physiological conditions.

Next, we characterized the secondary structures of five effective RNA fragments (T2, T3, T5, L4, L5) and two inactive RNA fragments (T8 and T9) under denaturing conditions using SHAPE-MaP (selective 2′-hydroxyl acylation analyzed by primer extension and mutational profiling)^16,30,31^. Intriguingly, each effective RNA fragment displayed a large internal multi-branched loop (> 34 nt in length), whereas the inactive RNAs lacked this feature (Supplementary Fig. S2a, b). These findings further indicate that stable but loosely organized RNA architectures are particularly effective at preventing TDP-43 condensation.

### Loosely structured multi-branched loop domains are essential for preventing TDP-43 condensation

We next sought to determine whether loosely structured RNA domains are required to disrupt TDP-43 condensation. The 332-nt T3 RNA, the smallest effective RNA fragment identified in the initial screen, was selected as a representative model (Supplementary Fig. S2a). T3 contains an 83-nt multi-branched loop^32^ with multiple UG-repeats (Motif 3) and three duplex-containing regions (Motifs 1, 2, and 4) (Fig. 1e). Deletion of Motif 3 completely abolished the ability of T3 to inhibit TDP-43 condensation *in vitro*, while individual or combinational deletion of the three duplex-containing regions had no detectable effect (Fig. 1f). These results indicate that the large multi-branched loop was indispensable for T3 RNA’s anti-TDP-43 condensation activity. Notably, deletion of Motif 1 modestly reduced T3 activity (relative turbidity ∼0.5; Fig. 1f), likely because this structure was formed by base-pairing between the terminal regions of T3 and may help maintain the overall conformation of the RNA.

To further validate the functional importance of such multi-branched loop structures in preventing TDP-43 condensation, we examined the denatured L4 RNA as a second example. It comprised two otherwise inactive fragments (T8 and T12) that flank a 50-nt multi-branched loop structure (L4-M) (Supplementary Fig. S3a), suggesting that L4-M might underlie the anti-condensation activity of the full-length L4. Indeed, while the isolated L4-M RNA itself showed no inhibitory effect, a stabilized variant (L4-ML) containing an extended 7-bp stem displayed detectable activity (Supplementary Fig. S3a-c). These results indicate that both the presence and structural stability of the multi-branched loop influence anti-condensation function.

Given the considerable length of L4 RNA (483 nt), we designed a series of truncations to identify the structural elements required for activity. In addition to the multi-branched loop, L4 contains a high-affinity TDP-43-binding motif (TDP-43 BM) and a terminal hairpin structure (EHS) analogous to the reported neuroprotective *Clip34* RNA via targeting TDP-43 toxicity^33^ (Supplementary Fig. S3b). To test whether the extended terminal pairing could stabilize the multi-branched loop, we included a linker sequence (LS) in certain variants. A panel of eight truncated RNAs (L4-1 to L4-8) was generated by combining these elements in different configurations (Supplementary Fig. S3d).

Turbidity assays showed that the activity of L4 RNA depended primarily on its loosely organized multi-branched loop, whereas additional stabilizing or TDP-43-binding elements provided limited and context-dependent benefits. Specifically, extended base-pairing did not enhance, but in some cases reduced, anti-TDP-43 condensation activity. Inclusion of either the TDP-43 BM or EHS alone also failed to improve efficacy. Only when both elements were combined, a moderate enhancement was observed (Supplementary Fig. S3e). Among the variants, L4-7 exhibited the strongest activity, comparable to full-length L4, and was selected for further investigation. Similar to what we have observed for T3, the multi-branched loop in L4-7 contained several duplex-containing substructures (Motifs 1-3; Supplementary Fig. S3f), and deletion of these substructures did not impair the ability of L4-7 to prevent TDP-43 condensation (Supplementary Fig. S3g).

Collectively, these data strongly suggest that loosely structured multi-branched loops, rather than stable duplexes or isolated high-affinity binding motif, are essential for suppressing TDP-43 condensation.

### Circular RNA aptamers outperform their linear counterparts in suppressing TDP-43 condensation

Circular RNAs possess elevated conformational stability via their covalently closed topology, we therefore hypothesized that circularizing these multi-branched loop-containing RNA fragments may improve their ability to suppress TDP-43 condensation, relative to their potentially base-paired linear counterparts (Fig. 1e). Consistent with this hypothesis, at an RNA: TDP-43 molar ratio of 1:10, although circularized T3 (circT3; Fig. 1g) and linear T3 showed comparable, strong activity, the circularized L4-ML (circL4-ML) outperformed its linear form (L4-ML) (Supplementary Fig. S3h). Based on its overall efficacy, circT3 was selected as the lead Capt candidate (Supplementary Fig. S3h).

To generate a more compact functional aptamer, we circularized the core 89-nt multi-branched loop motif of T3 (Motif 3), producing the 116-nt circT3-M3 (Fig. 1g). circT3-M3 retained activity comparable to circT3 and was approximately ∼2-fold more active than its linear counterpart (Fig. 1h). To explore whether activity could be further enhanced, we generated circT3-M3 variants containing either a TDP-43-binding motif (circT3-M3-BM) or a core binding sequence (circT3-M3-CBS) (Supplementary Fig. S3i). Neither variant improved activity relative to circT3-M3 (Fig. 1h), indicating that an additional binding element outside the central loop does not enhance anti-TDP-43 condensation activity.

Note that a circularized poly(A) control (circPoly(A)), containing a large and stable loop, showed no inhibition of TDP-43 condensation (Supplementary Fig. S3j, k). This confirms that sequence context, such as the UG repeats in circT3 and circT3-M3, rather than loop topology alone, is also important for its inhibitory role on TDP-43 condensation.

### Circular RNA aptamers suppress TDP-43 condensation sub-stoichiometrically and in a time-dependent manner

We next evaluated the activity of circT3 and circT3-M3 at low stoichiometric ratios via turbidity assay. Remarkably, both Capts suppressed TDP-43 condensation at RNA: protein ratios as low as 1:1,280, whereas linear T3 lost activity at a 1:320 ratio (Fig. 1i). Furthermore, at the 1:100 molar ratio, this inhibitory effect started to be detectable after 2 min of incubation of TDP-43 with RNA aptamers and became significant by 5 min incubation (Fig. 1j).

To confirm this finding, we introduced an EGFP tag to the C-terminus of TDP-43 to make TDP-43-EGFP-TEV-MBP recombinant protein and applied it for imaging analysis. Following RNA aptamer incubation and TEV protease cleavage, we observed that TDP-43-EGFP alone formed solidified condensates, and such condensation was effectively inhibited by the RNA aptamers (Supplementary Fig. S4a). Notably, consistent with the turbidity results, circT3 and circT3-M3 outperformed linear T3 at low RNA concentrations and remained active at the RNA: protein ratio of 1:1,280 (Supplementary Fig. S4a, b). Furthermore, all tested RNA aptamers exhibited detectable activity starting at 2 min of incubation and acted in a time-dependent manner (Supplementary Fig. S4c, d). These results together show that such Capts can regulate TDP-43 at levels far below their own abundance, and that the time-dependent transition further suggests a chaperone-like mechanism, analogous to our previous described *SLERT*-DDX21 model^23^.

Finally, we compared the activity of these RNA aptamers with that of the reported *Clip34* and *Malat1_start*^13^. At an RNA: TDP-43 ratio of 1:10, we found that *Clip34* displayed very limited capacity to suppress TDP-43 condensation, whereas *Malat1_start* was completely ineffective. Both of them appeared much less potent than T3, circT3 and circT3-M3 (Supplementary Fig. S4e). Furthermore, combining *Clip34* or *Malat1_start* with our established RNA aptamers did not synergize, but rather attenuated inhibition of TDP-43 condensation, likely due to their competitive binding to TDP-43 (Supplementary Fig. S4e).

Together, we identified two Capts, circT3 and its compact derivative circT3-M3, that contain multi-branched loops, could suppress TDP-43 condensation *in vitro* at remarkable sub-stoichiometric levels, outperforming their linear counterparts (Fig. 1k).

### RNA aptamers bind key residues in RRM1 to maintain TDP-43 in a soluble oligomeric state

To understand how these RNA aptamers suppress TDP-43 condensation, we first mapped their binding sites on TDP-43. Electrophoretic mobility shift assays (EMSAs) revealed that all of the RNA aptamers bind preferentially to the RRM1 domain rather than to RRM2 at RNA: protein ratios as low as 1:5. Combining both RRMs did not further enhance binding affinity beyond that observed with RRM1 alone (Fig. 2a).

**Fig. 2.**
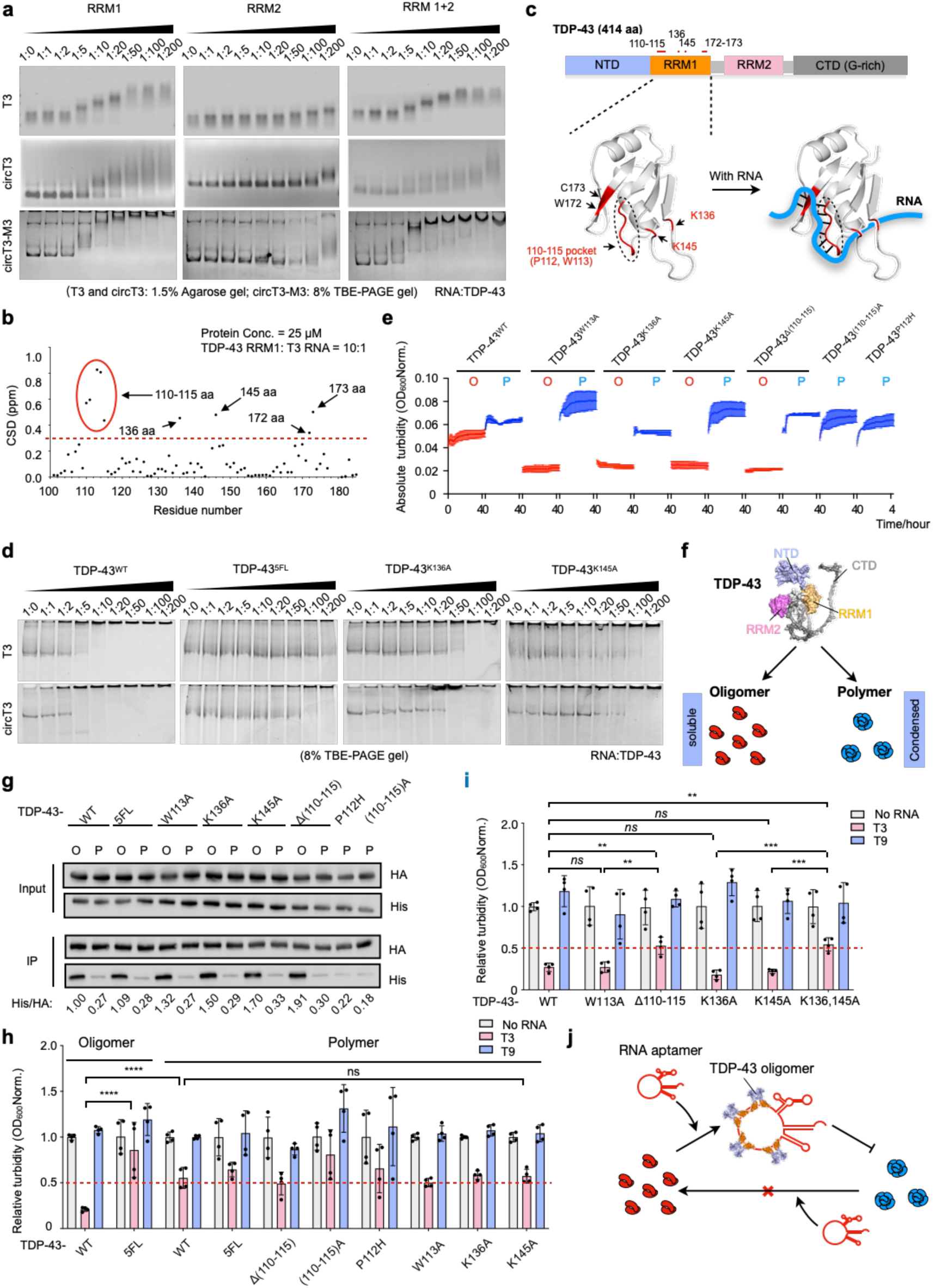
Key residues at TDP-43 RRM1 domain are required to maintain TDP-43 at soluble oligomeric state. **a** EMSA experiments show that the RNA aptamers have high affinity to the RRM1 domain, rather than to the RRM2 domain, of TDP-43. **b** The chemical shift differences (CSDs) of TDP-43 RRM1 domain with the addition of T3 RNA fragment, obtained by the nuclear magnetic resonance (NMR) spectroscopy at protein: RNA = 10:1. The amino acids that have the CSDs over 0.3 ppm are considered as binding sites to the RNAs. **c** Schematic to show T3 RNA binds the key residues at TDP-43 RRM1 domain. Upper: the domain architecture of TDP-43. Bottom: the structural schematic to show TDP-43 and T3 interaction, drawn with the help of Pymol software. The binding sites are marked in red. Reported functional and pathological residues are labeled in red and other residues are labeled in black. Interacting RNA is drawn in blue. **d** EMSA experiments show that T3 and circT3 RNA interact with MBP-tagged TDP-43^WT^ oligomers, but not MBP-tagged TDP-43^5FL^ oligomers. This interaction is impaired when K136 or K145 is mutated to alanine (K136A or K145A). **e** Turbidity detection of the oligomeric and polymeric MBP-tagged TDP-43 variants shows that TDP-43 oligomers are soluble, whereas TDP-43 polymers readily form condensates. **f** Schematic to show that TDP-43 has both soluble oligomeric and condensed polymeric states. **g** TDP-43 oligomers, but not polymers, show high homomeric affinity, revealed by the co-immunoprecipitation of the purified C-terminal MBP-HA-tagged and MBP-His-tagged TDP-43 oligomers (O) or polymers (P). **h** Turbidity assays reveal that the T3 RNA aptamer markedly inhibits the condensation of wild-type TDP-43 oligomers, yet exerted no significant effects on TDP-43^5FL^ oligomers or TDP-43 polymeric variants. T9 RNA is used as negative control. In this study, the oligomeric and polymeric MBP-tagged TDP-43 variants are pre-incubated with T3 or T9 RNA for 10 min. After that the mixed samples are treated with TEV enzyme for 15 min followed by turbidity detection. **i** Turbidity assays reveal that T3 RNA has relatively mild efficacy to prevent the condensation of TDP-43^△110–115^ and TDP-43^K136,145A^ oligomers compared to wild-type, W113A, K136A, and K146A TDP-43 variants. **j** Schematic to show that T3 RNA aptamer prevents the condensation of TDP-43. However, it has mild efficacy in dissolving the formed polymeric condensates.

We next used nuclear magnetic resonance (NMR) to identify T3-binding residues within RRM1. Addition of T3 RNA induced significant chemical shift deviations (CSDs > 0.3 ppm) at residues 110-115, 136, 145, 172, and 173 (Fig. 2b, c; Supplementary Fig. S5a), indicating these residues comprise the primary interaction interface. Notably, this mapped interface overlaps with residues previously implicated in RNA recognition, disease-associated variation, and ALS-linked acetylation. The 110-115 pocket is crucial for selectively binding UG-rich RNAs, with P112 and W113 serving as core binding sites^34^. P112H is a pathological variant associated with frontotemporal lobar degeneration (FTLD)^35^. In addition, K136 and K145 were identified as acetylation sites, with elevated acetylation levels reported in ALS patients^36,37^. Notably, mutating these residues, such as K136 or K145 to alanine, significantly inhibited the interaction between TDP-43 variants and T3 or circT3 RNAs (Fig. 2d). As a negative control, the TDP-43^5FL^ variant (F147, F149, F194, F231, F233 to L mutation; Supplementary Fig. S4a) that barely bound RNA^33^ exhibited almost no detectable interaction with either aptamer (Fig. 2d). Thus, our reported RNA aptamers engage a functionally and pathologically sensitive surface within RRM1.

It is known that TDP-43 exists either in native oligomeric state, consisting of a spectrum of species predominantly including dimers, or in LCD-driven pathological polymeric states^9,38^. Intriguingly, when purifying TDP-43 variants (Supplementary Fig. S5, b-d) and subjected them for turbidity assay, we found that oligomeric TDP-43 remained soluble, whereas polymeric TDP-43 readily formed condensates (Fig. 2e, f). In addition, the purified TDP-43 oligomers underwent progressive conversion into polymers following incubation (Supplementary Fig. S5e, f). To further validate that TDP-43 polymers, rather than oligomers, are the condensation-competent species, we purified a panel of oligomeric TDP-43 variants, including N- and C-terminal truncations and point mutants (S48E, 6M)^9^, that do not readily convert into polymers (Supplementary Fig. S6a, b)^9,39,40^. As expected, all these variants remained predominantly oligomeric after 2 h incubation at room temperature (RT) and remained soluble in turbidity assays (Supplementary Fig. S6c-f). Furthermore, treatment with T3 RNA did not substantially alter their condensation profiles (Supplementary Fig. S6f), indicating that aptamer activity is most apparent when TDP-43 retains the capacity to transition from an oligomeric state into pathological polymeric condensates.

Given the oligomeric state of TDP-43 is reported to be stabilized by the homomeric interactions through its NTD^9^, we compared homomeric interactions between oligomeric or polymeric TDP-43 variants carrying mutations in the RNA-interaction surface (Fig. 2c). Wild-type TDP-43 and TDP-43^5FL^ variant served as controls. Using C-terminal HA- and His-tagged TDP-43 variants under conditions favoring either oligomeric or polymeric states, we found that oligomeric TDP-43 engaged in stronger homomeric interactions than did polymeric forms (Fig. 2g). Of note, mutating the RNA interaction residues did not impair the interaction between TDP-43 oligomers (Fig. 2g).

Finally, we tested whether the RNA aptamers could reverse pre-formed TDP-43 polymeric condensates. As expected, the T3, but not T9, effectively suppressed condensation of wild-type oligomeric TDP-43. This inhibitory capacity was substantially diminished to TDP-43^5FL^ oligomer that lost RNA-binding capacity (Fig. 2h). Nevertheless, T3 RNA showed markedly reduced efficacy against the pre-formed TDP-43 polymeric condensates, including polymeric variants carrying mutations at RNA-interacting residues (Fig. 2h). Of note, we found that T3 RNA also reduced its capability to prevent condensation of oligomeric TDP-43^Δ110–115^ and TDP-43^136,145A^ mutants, likely due to the lack of binding to these proteins (Fig. 1i).

Collectively, these results indicate that these RNA aptamers primarily inhibit conversion of soluble oligomers into pathological polymers but are unlikely capable of reversing already-formed polymeric condensates (Fig. 2j).

### Circular RNA aptamers promote RRM1-mediated TDP-43 oligomerization

Of an important note, when purifying TDP-43 variants carrying mutations at RNA-interaction residues, we found that introducing the P112H mutation or substituting alanine into the 110-115 pocket completely abrogated the oligomeric state of TDP-43 (Supplementary Fig. S5c), suggesting that these residues at RRM1 may also contribute to TDP-43 homomeric assembly, in addition to the reported NTD^9^. This finding promoted us to investigate whether additional TDP-43 domains were also engaged for its homomeric interaction. Using AlphaFold3 to predict homomeric interfaces, we identified potential interaction sites within RRM1, RRM2, and the inter-RRM linker, in addition to the established N-terminal interaction region^9^ (Fig. 3a). Notably, K145, a residue critical for T3 RNA binding (Fig. 2c), also mapped to a predicted oligomerization interface (Fig. 3a), suggesting that RNA binding and oligomerization may converge on overlapping structural surfaces. In addition, 110-115 residues at RRM1 were not predicted as homomeric interaction sites (Fig. 3a), indicating their contribution to homomeric interaction was indirect.

**Fig. 3.**
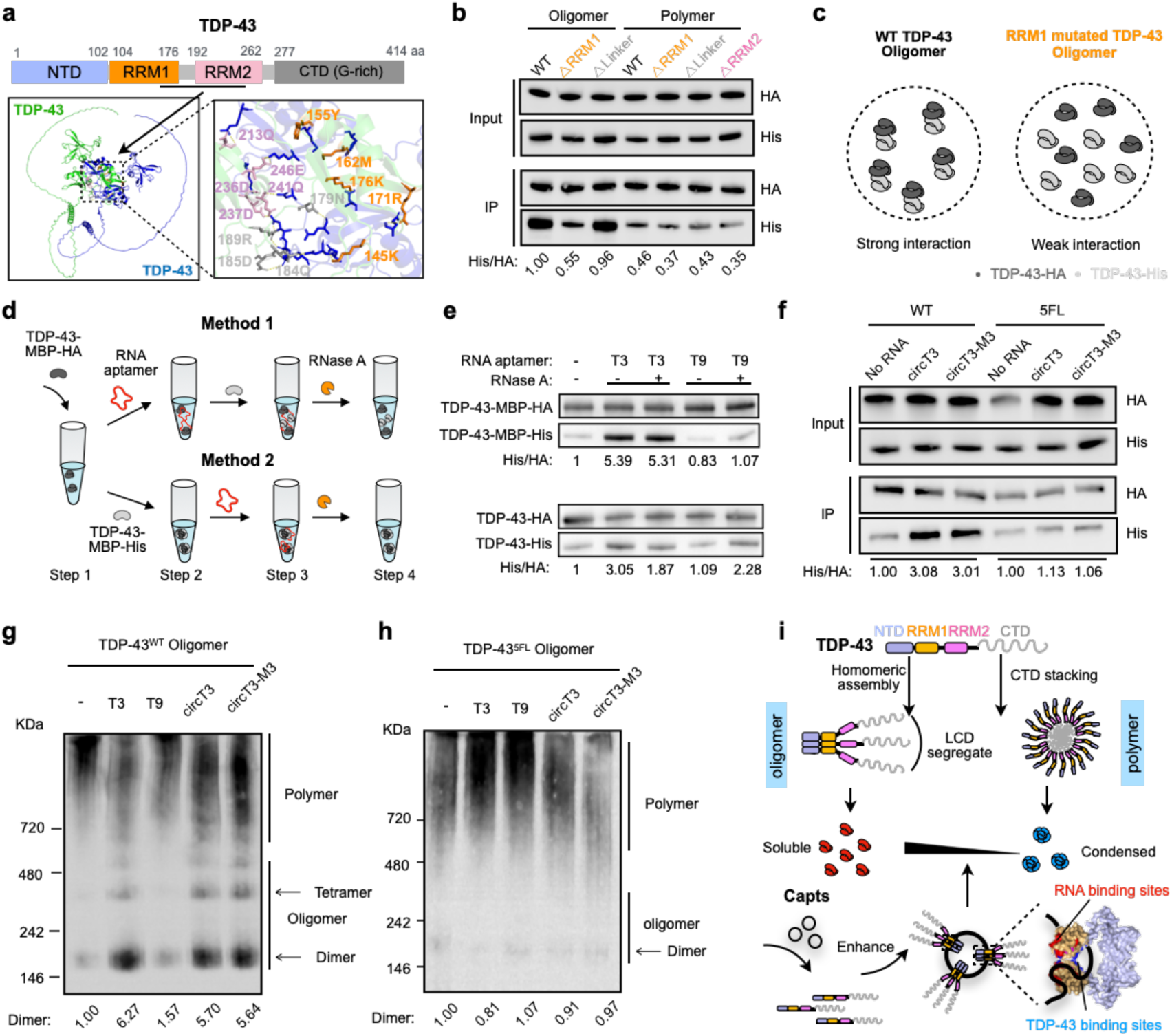
Circular RNA aptamers promote TDP-43 oligomerization by enhancing homomeric interactions. **a** Alphafold3 prediction of TDP-43 homomeric interaction. The interaction sites in RRM1 domain are marked in orange, in RRM2 domain are marked in purple, in the linker sequence between RRM1 and RRM2 are marked in grey. The interaction sites from the other interacting TDP-43 are marked in blue. **b** RRM1 mutated TDP-43 oligomers, but not wild-type or linker mutated TDP-43 oligomers, show impaired homomeric interaction. All the polymeric TDP-43 variants show weak homomeric interaction, revealed by co-immunoprecipitation of the purified C-terminal MBP-HA-tagged and MBP-His-tagged TDP-43 oligomers or polymers. **c** Schematic to show that RRM1 mutated TDP-43 oligomers have weaker homomeric interaction compared to wild-type TDP-43 oligomers. **d** Schematic for testing RNA aptamers in affecting TDP-43 homological interaction. Renatured RNA aptamers are added before (Method 1) or after (Method 2) TDP-43-MBP-HA and TDP-43-MBP-His protein incubation. The RNase A is added to remove RNA aptamers after incubation. **e** Addition of T3 but not T9 RNA (negative control) before protein incubation (Method 1) enhances the homomeric interaction between oligomeric TDP-43-MBP-HA and TDP-43-MBP-His. Addition of RNase A to remove the RNAs post incubation has mild effect on the interaction (upper). While addition of the RNA aptamers after protein incubation (Method 2) shows moderate impact to TDP-43 homomeric interaction (bottom), revealed by co-immunoprecipitation of the purified TDP-43. **f** The Capts promote homomeric interaction of oligomeric wild-type TDP-43, but not oligomeric TDP-43^5FL^ variant, revealed by the co-immunoprecipitation between the purified C-terminal HA-tagged MBP-TDP-43 and His-tagged MBP-TDP-43 variants. **g** T3, circT3, and circT3-M3 exert stronger inhibitory effects on TDP-43 polymerization than T9 RNA, revealed by the native-PAGE. In this study, wild-type MBP-tagged TDP-43 oligomers are incubated with different renatured RNA aptamers for 2h at RT, respectively, and then the samples are subjected for native-PAGE analysis. **h** T3, circT3, and circT3-M3 fail to inhibit the polymerization of MBP-tagged TDP-43^5FL^ oligomers, revealed by the native-PAGE analysis. **i** Schematic to demonstrate that TDP-43 either forms condensed polymers via the CTD stacking or forms soluble oligomers via the N-terminal and RRM1-mediated homomeric assembly. The oligomerization of TDP-43 spatially segregates the LCD of TDP-43, thereby preventing its polymerization. The Capts bind the key residues (red) at the RRM1 domain of multiple TDP-43 molecules to catalyze the RRM1-mediated (via the blue residues) TDP-43 homomeric assembly, thereby maintaining TDP-43 at the soluble oligomeric state.

We then purified both oligomeric and polymeric forms of TDP-43 variants carrying alanine mutations in these predicted regions (RRM1 mutation carrying Y155, M162, R171, and K176 to A; RRM2 mutation carrying Q213, D236, D237, Q241, and E246 to A; the linker region mutation carrying N179, Q184, D185, and R189 to A) (Supplementary Fig. S7a-d). Mutations in the RRM2 interface completely abolished oligomer formation, yielding only polymers (Supplementary Fig. S7d). In the oligomeric state, RRM1 mutations reduced homomeric interaction, whereas linker mutations had little effect (Fig. 3b, c). As expected, polymeric forms of all TDP-43 variants showed weaker homomeric interaction than wild-type oligomers (Fig. 3b, c).

We next asked whether these RNA aptamers could actively modulate TDP-43 homomeric interactions. Pre-incubation of HA-tagged TDP-43 oligomers with T3 RNA, but not the negative-control T9 RNA, significantly enhanced subsequent pulldown of His-tagged TDP-43 oligomers (Fig. 3d, e). Notably, RNase A treatment failed to abrogate this T3 RNA-mediated enhancement (Fig. 3e), suggesting that the RNA aptamers were not continuously required once the homomeric interaction had formed. This finding indicated that the RNA aptamers likely catalyze or guide TDP-43 oligomerization rather than functioning as a molecule glue. Consistent with this model, addition of RNA aptamers after TDP-43 proteins had already been incubated together produced only a moderate effect on homomeric interaction (Fig. 3e). As expected, circT3 and circT3-M3 aptamers similarly promoted homomeric interaction between wild-type TDP-43 oligomers, but not between RNA-binding-deficient TDP-43^5FL^ oligomers (Fig. 3f).

Furthermore, we evaluated the impact of RNA aptamers on TDP-43 assembly via native-PAGE analysis (Fig. 3g). The results showed that all active RNAs (T3, circT3, circT3-M3), but not the negative T9 RNA, reduced conversion of wild-type TDP-43 oligomers into higher-order polymers, preserving dimeric and tetrameric species (Fig. 3g). By contrast, these RNA aptamers failed to inhibit the polymerization of TDP-43^5FL^ oligomers that lack RNA binding capacity (Fig. 3h).

Together, our results suggest that the loosely structured T3 RNA, its circular form, and its minimized circular derivative can enhance RRM1-mediated homomeric assembly of TDP-43 to form soluble oligomers (Fig. 3e–g) by engaging key residues within RRM1 (Fig. 2c). This homomeric interaction likely spatially segregates the C-terminal LCD of TDP-43⁹, preventing its stacking and thereby inhibiting the formation of pathological polymers (Fig. 3i). This mechanism points to a potential route for therapeutic intervention in TDP-43-associated neurodegenerative disorders with such effective RNA aptamers.

### RNA aptamers chaperone TDP-43 at superb sub-stoichiometric ratios

Next, we asked how these RNA aptamers suppress TDP-43 condensation at such low stoichiometry. Inspired by the defining features of molecular chaperones^23,41^, we reasoned that an effective RNA chaperone should exhibit two key properties: the ability to rapidly bind multiple proteins and the capacity for reversible interaction. These properties enable one RNA molecule to sequentially engage and regulate numerous protein targets. Although the comparable size of an RNA and its binding partner may limit simultaneous occupancy, reversible association could enable superb sub-stoichiometric activity.

We employed single-molecule total internal reflection fluorescence (smTIRF) microscopy^19,23,42^ to directly assess the dynamic interactions between TDP-43 and these RNA aptamers. Biotin- and Cy5-labeled T3, circT3, and circT3-M3 were individually immobilized on a PEG-biotin-coated quartz surface via streptavidin, with similarly labeled T9 RNA serving as a negative control. Cy3-labeled TDP-43-MBP was then introduced to record binding events in real-time (Fig. 4a).

**Fig. 4.**
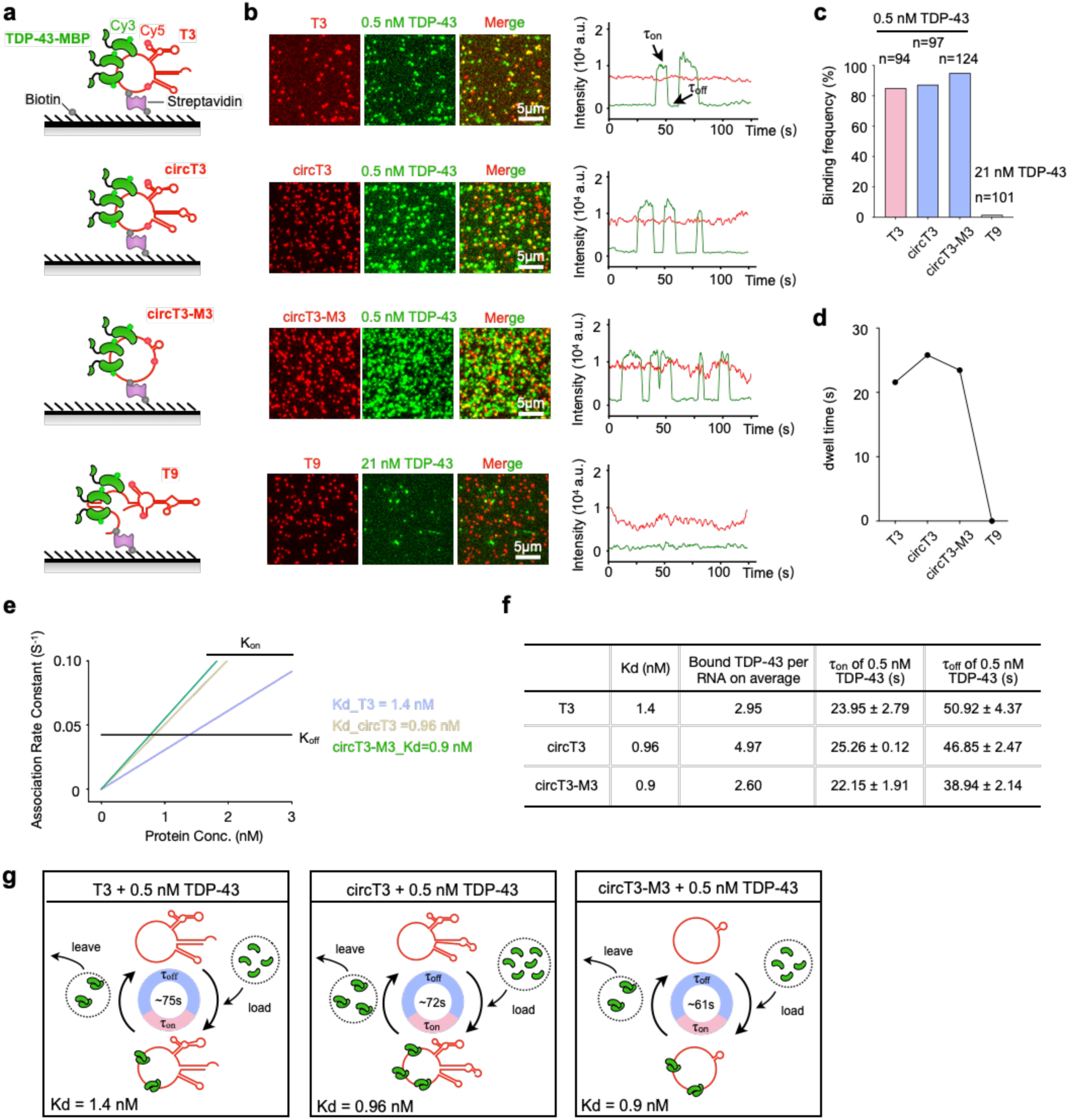
circT3 and circT3-M3 chaperone TDP-43 at a superb sub-stoichiometric ratio. **a** Schematic of smTIRF to image direct interactions between MBP-Cy3-tagged TDP-43 oligomers and biotin and Cy5-labeled T3, circT3, and circT3-M3 RNAs. Cy5 and biotin are internally labeled during IVT. After TDP-43-MBP-Cy3 injection, the single-molecule videos in two corresponding channels are recorded. The binding time (τ_on_) and disassociating time (τ_off_) are pointed by the arrowhead. T9 RNA is used as negative control. **b** T3, circT3, and circT3-M3, but not T9 interact with MBP-tagged TDP-43 oligomers, visualized by smTIRF. **c** Statistic analysis of the interaction between RNA aptamers and TDP-43 from **b**. 0.5 nM T3/circT3/circT3-M3 and 21 nM T9 are applied individually to capture the interacting TDP-43. **d** The dwell time analysis of MBP-tagged TDP-43 oligomers residing on T3, circT3, circT3-M3, and T9 from the smTIRF experiments. **e** The apparent dissociation constant (Kd) calculation between MBP-tagged TDP-43 oligomers and T3, circT3, or circT3-M3 interaction. K_on_ (association rate constant), K_off_ (dissociation rate constant). **f** The table documenting the interaction characteristics between RNA aptamers and MBP-tagged TDP-43 oligomers, including the Kd, bound TDP-43 molecules per RNA on average, and the association time (τ_on_) and dissociation time (τ_off_) for the RNA aptamers to interact with 0.5 nM TDP-43. **g** Schematic to show that the RNA aptamers transiently and iteratively interact with multiple TDP-43 molecules, acting as molecular chaperones to promote TDP-43 oligomerization.

Upon injection of 0.5 nM TDP-43, robust interactions with the active RNA aptamers were readily detected. In contrast, no binding was detected between T9 RNA and TDP-43, even at a protein concentration as high as 21 nM (Fig. 4b, c). Titrating experiment showed that individual TDP-43 molecule remained bound to T3 RNA for an average dwell time of ∼22 sec before dissociation (Fig. 4d; Supplementary Fig. S8a, b), yielding an apparent dissociation constant (Kd) of 1.4 nM (Fig. 4e). Under saturating protein conditions, photobleaching assays quantified that a single T3 RNA could simultaneously engage ∼3 TDP-43 molecules on average (Fig. 4f; Supplementary Fig. S8c). These findings suggest that T3 could act as an RNA chaperone, regulating TDP-43 molecules in excess of its own abundance.

We next compared the chaperone activity of Capts and Lapts to understand why circT3 and circT3-M3 outperformed linear T3 under low stoichiometry conditions (Fig. 1i). Although the apparent affinity (∼1 nM) and bound-state dwell time (∼25 sec) for TDP-43-Capts interaction were comparable to those of linear T3 (Fig. 4d, e; Supplementary Fig. S8a, b), circT3 exhibited a greater number of simultaneously bound TDP-43 molecules (Fig. 4f; Supplementary Fig. S8d). Meanwhile, circT3-M3 displayed the shortest binding-dissociation cycling time, despite binding fewer TDP-43 molecules, likely due to its reduced size (Fig. 4f; Supplementary Fig. S8e). The nM-level apparent binding kinetics between RNA aptamers and TDP-43 were further confirmed via titrated biolayer interferometry (BLI), a label-free platform that records real-time association and dissociation profiles (Supplementary Fig. S8f). Together, these enhanced binding characteristics collectively contribute to the super inhibitory efficacy of the Capts in suppressing TDP-43 condensation at low stoichiometry.

Collectively, these results (Fig. 4g) showed that our reported RNA aptamers could rapidly and reversibly interact with multiple TDP-43 molecules within a time cycle of 60-70 sec (Fig. 4f). This on-off-cycle interaction likely enabled them to act as a mode of RNA chaperone to catalyze multiple TDP-43 oligomerization efficiently, thereby achieving a low stoichiometric effect in a time-dependent manner (Fig. 1i, j; Supplementary Fig. S4a-4d). Of note, the Capts either engage more TDP-43 molecules (circT3) or carry a shorter cycling time (circT3-M3), thus offering the circular format RNA aptamers an augmented capacity to outperform the linear T3 (Fig. 4g).

### Circular RNA aptamers suppress intracellular TDP-43 condensation and restore TDP-43-dependent splicing defects

Prompted by the superb sub-stoichiometric activity of these Capts against TDP-43 condensation observed *in vitro*, we next asked whether they exhibit cellular efficacy superior to that of linear T3. To answer this question, we established SH-SY5Y and U2OS cell models exhibiting ectopic cytosolic TDP-43 condensation^43,44^. Cells were transfected with plasmids encoding mNG-tagged TDP-43 harboring a nuclear localization signal (NLS) mutation (mNG-TDP-43^NLS^), which localized predominantly to the cytoplasm, in contrast to nuclear localized mNG-TDP-43 (Supplementary Fig. S9a, b). Next, the transfected cells were stressed with 250 μM sodium arsenate (SA) for 1 h to induce condensation (Supplementary Fig. S9c)^43,44^. Under such condition, 48.7% of SH-SY5Y cells and 62.7% of U2OS cells displayed visible cytosolic mNG-TDP-43^NLS^ condensates, respectively (Supplementary Fig. S9d).

Before assessing their anti-TDP-43 condensation activity, we evaluated the intracellular stability of these RNA aptamers and their impact on native nuclear TDP-43 functions in wild-type SH-SY5Y cells. Both circT3 and circT3-M3 exhibited a half-life of ∼12 h, whereas linear T3 displayed a shorter half-life of less than 6 h (Supplementary Fig. S9e). RNA-seq revealed that delivery of both Capts induced only minimal alterations in global gene expression (Supplementary Fig. S9f) and alternative splicing (AS) events (Supplementary Fig. S9g, h), indicating that the function of endogenous TDP-43 was unlikely affected. Of note, these results were as expected, given the lipid nanoparticle (LNP)-delivered RNA were mainly localized to the cytosol^45^, whereas endogenous TDP-43 and their targets were in the nucleus.

Next, we delivered these RNA aptamers via LNP prior to SA stress (Fig. 5a; Supplementary Fig. S10a). To determine the minimum effective aptamer concentration, we treated cells with a concentration gradient of each aptamer (0.05 nM, 0.2 nM, 0.5 nM, 2 nM, and 5 nM). Imaging confirmed that all delivered RNA aptamers localized primarily to the cytoplasm in both cell lines (Fig. 5b; Supplementary Fig. S10b). In U2OS cells, linear T3 RNA showed detectable suppression of TDP-43 condensation starting at 0.5 nM, whereas an effect was observed at 2 nM in SH-SY5Y cells. In contrast, both Capts displayed potent activity at 0.2 nM in either cell line. None of the aptamers showed detectable activity at 0.05 nM (Fig. 5c-e; Supplementary Fig. S10c-e). Within the low-concentration range of 0.2-2 nM, Capts consistently outperformed linear T3 in suppressing TDP-43 condensation (Fig. 5c-e; Supplementary Fig. S10c-e). Of note, administering the aptamers post SA stress (Supplementary Fig. S10f) barely showed efficacy at either low concentration of 0.5 nM or high concentration of 5 nM (Supplementary Fig. S10g, h), which was consistent with our turbidity results that T3 RNA exhibited only modest activity in dissolving purified TDP-43 polymers (Fig. 2h). Collectively, these results suggest that circT3 and circT3-M3 can inhibit TDP-43 condensation at nM-scale, consistent with a chaperone-like, superb sub-stoichiometric mechanism, even within cells.

**Fig. 5.**
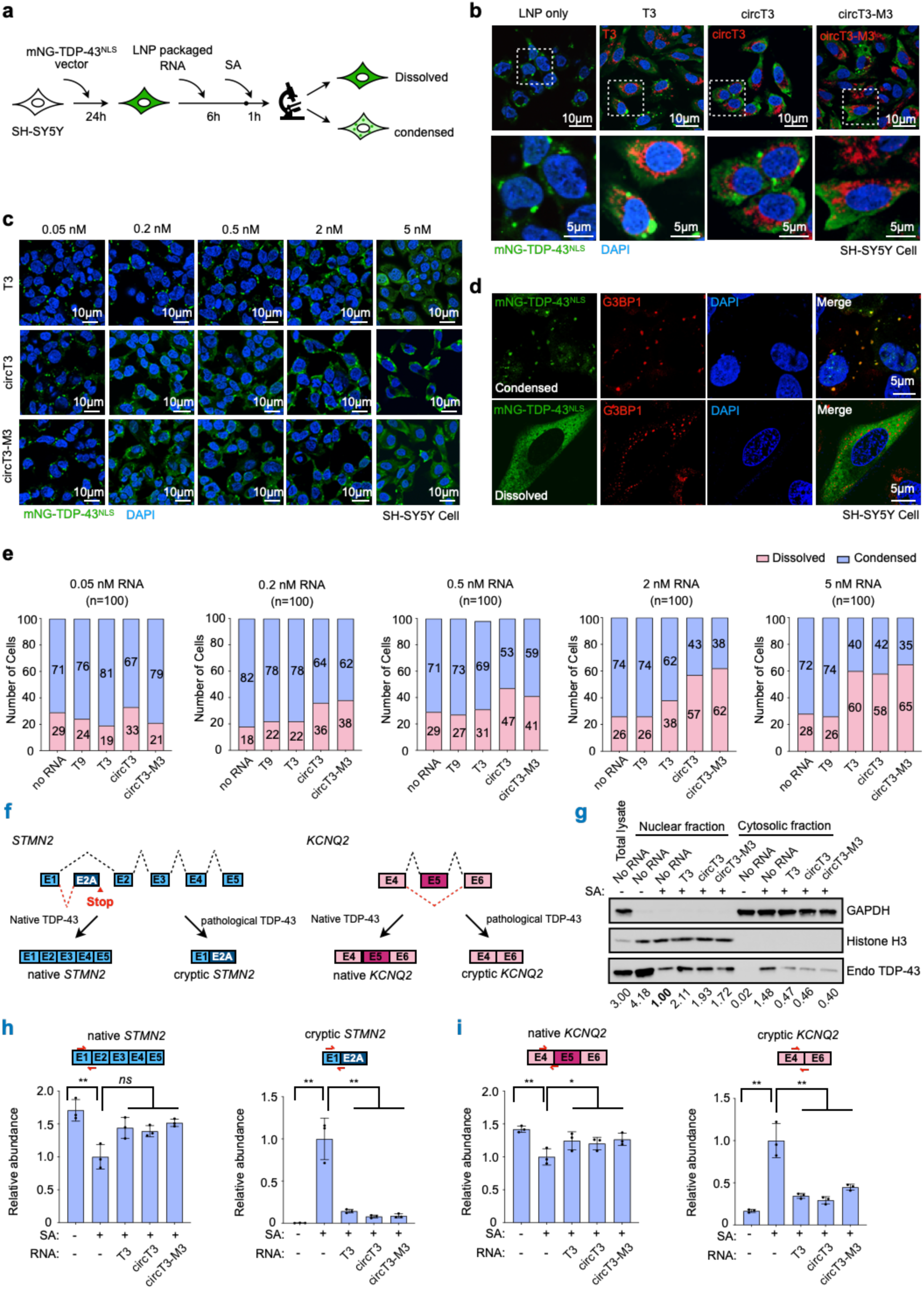
Circular RNA aptamers suppress TDP-43 condensation and restore TDP-43-dependent splicing defects in SH-SY5Y cells. **a** Schematic of employing LNP-packaged RNA aptamers to suppress TDP-43 cytosolic condensation in the SH-SY5Y cell model. **b** Representative images showing the delivered RNA aptamers in the SH-SY5Y cell model. mNG-TDP-43 (green), RNA aptamers (red). The nucleus is stained with DAPI. Scale bar = 5 μm. **c** Representative images showing the condensation state of TDP-43 post RNA aptamer delivery at concentrations of 0.05 nM, 0.2 nM, 0.5 nM, 2 nM, and 5 nM in the SH-SY5Y cell model. mNG-TDP-43 is stained in green. The nucleus is stained with DAPI. Scale bar = 10 μm. **d** Representative zoom-in images showing the aggregated (upper) and dissolved (bottom) TDP-43 post RNA aptamer delivery in the SH-SY5Y cell model. mNG-TDP-43 (green), G3BP1 (red). The nucleus is stained with DAPI. Scale bar = 5 μm. **e** Statistics from (**c**) showing that the Capts exhibit inhibitory efficacy to TDP-43 condensation start from the concentration of 0.2 nM, outperforming T3 which initiates efficacy that start from 2 nM. **f** Schematic showing the TDP-43-dependent splicing and cryptic splicing of *STMN2* and *KCNQ2*. **g** WB analysis of nuclear and cytosolic fractions from the established SH-SY5Y cell model reveals that T3, circT3, and circT3-M3 moderately restore nuclear localization of endogenous TDP-43 following SA treatment. **h, i** qRT-PCR reveals that addition of RNA aptamers substantially rescues the mis-splicing of STMN2 (**h**) and KCNQ2 (**i**).

Loss of TDP-43 was known to induce cryptic splicing events in *STMN2* and *KCNQ2*, which contributes to the pathological manifestations of TDP-43 proteinopathy^46,47^ (Fig. 5f). Importantly, our characterized RNA aptamers could alleviate TDP-43-dependent aberrant splicing of *STMN2* and *KCNQ2* in the SH-SY5Y cellular model harboring cytosolic TDP-43 condensation. First, we observed that RNA aptamers treatment moderately restored nuclear localization of endogenous TDP-43 following SA treatment (Fig. 5g). Second, cryptic splicing of *STMN2* and *KCNQ2* detected in these cells could be substantially rescued by these aptamers (Fig. 5h, i).

As the immunogenicity of delivered RNA aptamers is one key feature for future application, we also assessed to what extent these aptamers could induce inflammatory gene expression in cells. Compared with poly(I:C) stimulation, linear T3 RNA elicited a moderate inflammatory response, as indicated by the elevated TNF-α, IFN-β, and IL-6 levels (Supplementary Fig. S10i). In contrast, both Capts barely induced the expression of these cytokines, confirming their low immunogenicity as circular RNAs (Supplementary Fig. S10i), consistent with previous reports^18,19^.

### Circular RNA aptamers reduce cytosolic TDP-43 accumulation and partially ameliorate TDP-43 proteinopathy-driven motor dysfunction *in vivo*

Lastly, we evaluated whether circT3 and circT3-M3 are effective *in vivo* by using the NEFH-tTA/tetO-hTDP-43^ΔNLS^ mouse model (Fig. 6a). This model enables doxycycline (Dox)-suppressible expression of human TDP-43 (hTDP-43) carrying a defective NLS (ΔNLS) under the *neurofilament heavy chain* (*NEFH*) promoter^48^. Upon Dox withdrawal, these mice develop progressive TDP-43 proteinopathy in nerves systems^48^.

**Fig. 6.**
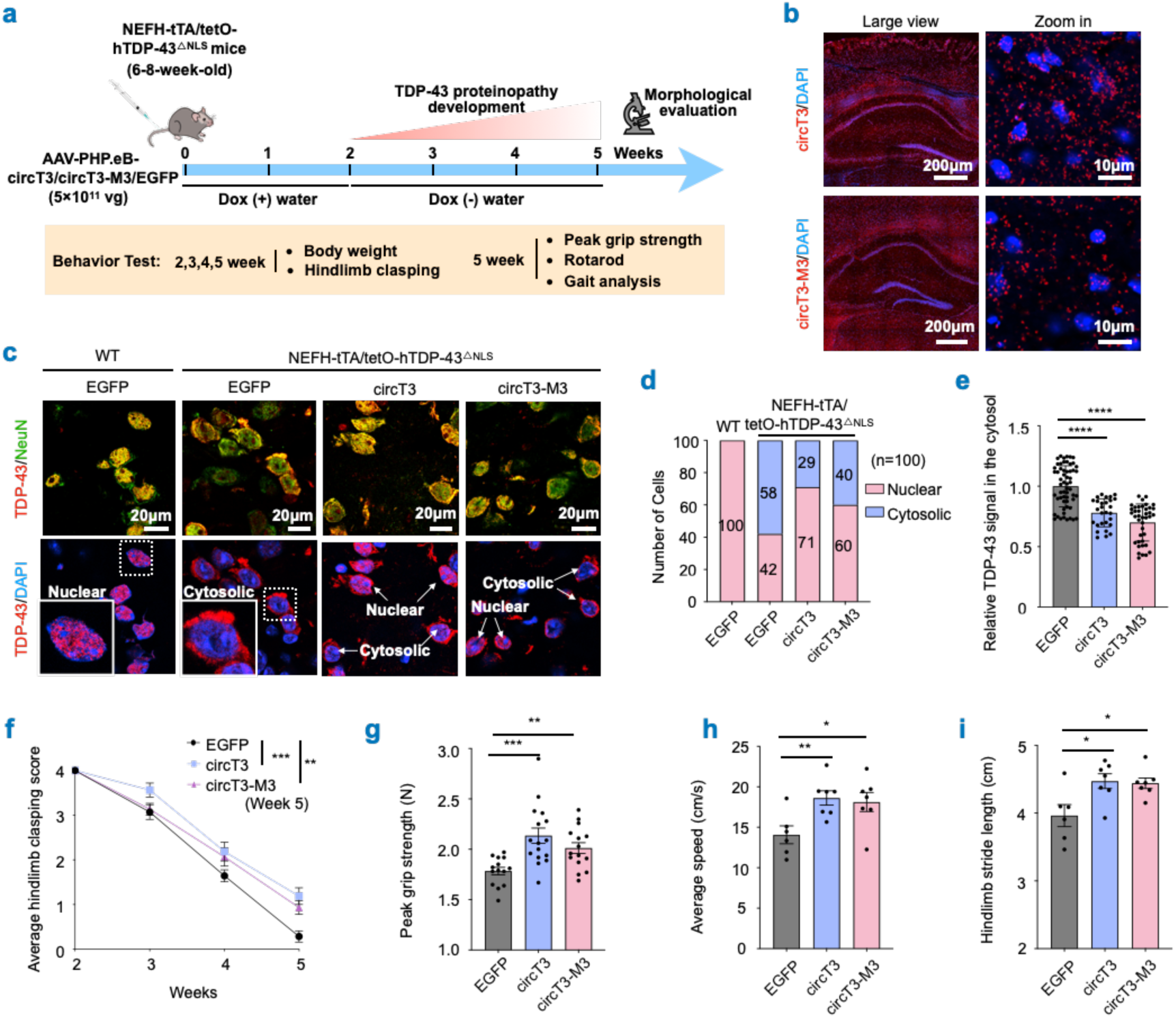
Circular RNA aptamers ameliorate TDP-43 proteinopathy-driven phenotypes in the mouse model. **a** Schematic showing that 6-8-week-old NEFH-tTA/tetO-hTDP-43^ΔNLS^ mice are administered AAV-PHP.eB-packaged plasmids encoding circT3, circT3-M3, or EGFP via tail-vein injection. Two weeks post-AAV injection, Dox is withdrawn to trigger TDP-43 proteinopathy in these mice. Morphological and behavioral tests are performed at distinct time points. **b** Representative large view and zoom in images showing the expression of circT3 or circT3-M3 in mouse brain sections. The nucleus is stained with DAPI. Scale bar = 200 μm (large view) and 10 μm (zoom in). **c** Representative images showing the expression pattern of TDP-43 (red) in mouse brain sections. The nucleus is stained with DAPI. Scale bar = 20 μm. **d** Statistics from (**c**) showing that delivery of AAV-PHP.eB-circT3 or AAV-PHP.eB-circT3-M3 reduces the cell numbers displaying TDP-43 cytosolic accumulation in NEFH-tTA/tetO-hTDP-43^△NLS^ mouse model compared to AAV-PHP.eB-EGFP control. n=100 for each group. **e** Statistics from (**c**) showing that cells with cytosolic TDP-43 accumulation display reduced TDP-43 signal intensity in Capts-treated groups relative to EGFP group. **f** Averaged hindlimb clasping scores from AAV-treated NEFH-tTA/tetO-hTDP-43^ΔNLS^ mice show that the Capt expression partially ameliorates hindlimb clasping deficits at week 5. **g** Peak grip strength measurements from AAV-treated NEFH-tTA/tetO-hTDP-43^ΔNLS^ mice show that the Capt expression improves muscle strength at week 5. **h-i** Gait analysis reveals that Capt-treated mice display improved running speed (**h**) and hindlimb stride length (**i**), compared to EGFP-expressing mice.

We administered AAV-PHP.eB-packaged plasmids encoding circT3 or circT3-M3 via tail-vein injection^49^ when the mice reached 6-8 weeks of age (Fig. 6a). Two weeks after that (week 2), Capts expression was detected in mouse brain sections (Fig. 6b), and Dox was then withdrawn. Mice injected with AAV-PHP.eB-packaged EGFP-expressing plasmids served as controls. Three weeks after Dox withdrawal (week 5), we observed cytosolic TDP-43 accumulation in ∼58% of the NeuN-positive neuronal cells in brain from NEFH-tTA/tetO-hTDP-43^ΔNLS^ mice injected with AAV-PHP.eB-EGFP, whereas all TDP-43 signals remained nuclear in their wild-type littermates (Fig. 6c, d). Furthermore, NEFH-tTA/tetO-hTDP-43^△NLS^ mice receiving AAV-PHP.eB-circT3 or AAV-PHP.eB-circT3-M3 harbored fewer NeuN-positive neuronal cells (∼29% and 40%, respectively) displaying cytosolic TDP-43 accumulation (Fig. 6c, d). Among cells with detectable cytosolic TDP-43 signals, groups expressing Capts exhibited ∼20-30% lower signal intensity relative to the EGFP-expressing group (Fig. 6e), suggesting that these Capts could mitigate TDP-43 cytosolic accumulation *in vivo*.

We next assessed whether these Capts could ameliorate pathological phenotypes triggered by TDP-43 proteinopathy in this mouse model (Fig. 6a). We monitored body weight and hindlimb clasping scores weekly post-AAV injection (Week 2-5). Mice receiving AAV-PHP.eB-EGFP developed body-weight loss starting at week 3, whereas delivery of AAV-PHP.eB-circT3 or AAV-PHP.eB-circT3-M3 failed to rescue this weight reduction (Supplementary Fig. S11a). However, Capts treatment ameliorated hindlimb motor dysfunction, as evidenced by improved average hindlimb clasping scores from 0.29 to 1.19 and 0.93, respectively, at week 5 (Fig. 6f). This finding inspired us to evaluate muscular strength and overall motor performance in these mice at this time point. First, we observed that mice receiving Capt treatment exhibited an approximately 10-20% increase in peak grip strength relative to EGFP-expressing control animals (Fig. 6g). Second, Capt-treated mice displayed a ∼30% improvement in running speed and a ∼12% increase in hindlimb stride length compared with EGFP-treated controls (Fig. 6h, i), although other higher-order gait parameters such as hindlimb duty cycle, hindlimb swing phase, diagonal support, and rotarod performance were not significantly improved (Supplementary Fig. S11b-e). Nonetheless, these results are inspiring that both circT3 and circT3-M3 can at least partially ameliorate TDP-43 proteinopathy-driven motor dysfunction *in vivo*.

## DISCUSSION

Many diseases are driven by pathological mislocalization, conformational conversion, or abnormal assembly of critical proteins, rather than by alterations in expression level alone^1,2,50^. In the nervous system, amyloid-β (Aβ) aggregation and tau hyperphosphorylation contribute to the progression of Alzheimer’s disease (AD)^51^, α-synuclein misfolding and aggregation are central to Parkinson’s disease (PD)^52^, and cytoplasmic mislocalization and aggregation of TDP-43 define most ALS case^4,5^. A major therapeutic challenge for these diseases is to selectively neutralize pathological protein states at the subcellular level while preserving normal protein function. Conventional strategies such as RNA interference or CRISPR-based depletion reduce total protein abundance and generally cannot distinguish physiological from pathological forms.

With covalently closed topology, circular RNAs exhibit greater aptamer potential than linear RNAs in terms of stability and structural rigidity^16,17,19,21^. Importantly, synthesized circular RNAs exhibit low immunogenicity, giving them great potential for RNA therapeutics^18,19^. Our recent work has shown that ds-cRNAs can inhibit aberrant PKR activation with high efficacy^16,20,21^. Here, we extend this concept to a proteinopathy context by identifying circT3 and its minimized derivative, circT3-M3, as Capts that suppress TDP-43 condensation. This finding indicates that Capts can be used not only to block aberrant enzyme activation, but also to redirect pathological protein assembly.

lncRNA can modulate target proteins at low stoichiometry^53^, but the mechanism is unclear. Among them, *NORAD* induces LLPS of Pumilio (PUM) to sequester excess PUM in NP bodies to modulate genome stability^54^. *SLERT*, at levels far below DDX21, chaperons DDX21 into a closed conformation, preserving nucleolar properties and RNA Polymerase I transcription^23^. Here, we show that our characterized Capts adopt a novel chaperone-like mechanism to suppress TDP-43 condensation in a superb sub-stoichiometric manner, achieving an RNA: protein molar ratio as low as approximately 1:1,000 (Fig. 1i, j; Supplementary Fig. S4a-4d). They employ single-stranded structures to chaperone several TDP-43 molecules at once through iterative binding events, catalyzing RRM1-mediated homomeric interactions among TDP-43 proteins. This process enhances N-terminus and RRM1-driven TDP-43 oligomerization, which effectively outcompetes the C-terminal prion-like domain-mediated LLPS and supersaturation (Fig. 2-4; Supplementary Fig. S5-S8). This distinct chaperone-like working model to suppress TDP-43 condensation distinguishes them from the mechanisms of ATTEC-directed mHTT degradation in Huntington’s disease and SPYTAC-directed Aβ clearance in Alzheimer’s disease^55,56^, neither mode of action of *NORAD*^54^ or *SLERT*^23^. Of note, RNase A treatment barely exerts an impact on the TDP-43 homomeric interaction once formed (Fig. 3d, e). Conversely, these aptamers show reduced efficacy in reversing pre-existing TDP-43 polymeric condensates (Fig. 2h; Supplementary Fig. S10f-10g), suggesting that they are best suited for preventing or slowing the deterioration of TDP-43-related proteinopathy rather than reversing late-stage diseases.

Of an important note, RNA structures appear to be essential to effectively prevent TDP-43 condensation. In the linear RNA context, denatured RNAs show higher efficacy compared with their renatured format (Fig. 1c, d). Given that TDP-43 prefers to bind single-stranded RNAs^28^, we propose that the denatured RNAs likely provide less-structured and single-stranded regions to effectively engage with TDP-43 molecules. In line with this, all tested positive RNAs have a reasonably large internal loop structures to enable the single-stranded region (Supplementary Fig. S2), such as the Motif 3 in T3 RNA (Fig. 1e). Consistently, circularization of such RNA fragments likely enables the formation of less-structured regions to be more stable, thereby offering an even more efficient molecular format to interact with TDP-43 (Supplementary Fig. S12). Additional experiments, such as the high-resolution structure analyses of these RNA aptamers and TDP-43-RNA complexes are warranted to further delineate their modes of action with TDP-43.

Finally, it is noteworthy that both Capts effectively suppress TDP-43 condensation in cellular models and restore TDP-43-dependent splicing defects at nM-level with minimal disruption to the endogenous nuclear TDP-43 activity and negligible immunogenicity (Fig. 5; Supplementary Fig. S9-S10). Importantly, these Capts partially ameliorate TDP-43 proteinopathy-driven motor dysfunction in the mouse model (Fig. 6), providing the first *in vivo* evidence that such neurodegenerative phenotypes can be remitted by RNA aptamers. Together, the superior subcellular targeting efficacy, regulatory activity at low doses, and minimal side-effects of Capts making them as promising therapeutic alternatives for TDP-43-associated proteinopathies. Given that many neurodegenerative disorders are driven by pathological protein alterations, this study may offer a conceptual framework for developing Capt-based therapeutic strategies applicable to broader spectrum of such diseases.

Several questions remain at this stage. First, our *in vitro* and in-cell experiments strongly suggest that the characterized Capts efficiently suppress TDP-43 condensation, but the exact mechanism for preventing TDP-43 aggregation requires further studies. Second, our *in vivo* experiments have shown that these aptamers can mitigate cytosolic TDP-43 accumulation and partially ameliorate TDP-43 proteinopathy-driven motor dysfunction. More comprehensive evaluation of nerve functional recovery, tracking of long-term efficacy, and safety should be further conducted. Third, we notice that the higher-order behavioral performance of these mice is not significantly improved by the Capts, indicating additional therapeutic strategies, such as specifically degrade the pathological forms of TDP-43, should be combined to fully address this disease.

## MATERIALS AND METHODS

### Experimental animals

All animal procedures were approved by the Institutional Animal Care and Use Committee of the Center for Excellence in Molecular Cell Science, Chinese Academy of Sciences, under an approved protocol. Mice were housed with littermates in a specific-pathogen-free facility and monitored routinely. NEFH-tTA line 8 and tetO-hTDP-43-ΔNLS line 4 mice were obtained from The Jackson Laboratory and crossed to generate NEFH-tTA/tetO-hTDP-43ΔNLS offsprings. Mice received Dox-containing drinking water (2 mg/mL) until Dox withdrawal two weeks after administration of AAV-PHP.eB carrying the plasmid expressing circT3, crcT3-M3, or EGFP (5×10^11^ vg in 200 µL PBS). Genotypes were determined from tail DNA using the primers listed in Table S2.

### Behavior experiments

Behavioral experiments were performed between 8:00 a.m. and 8:00 p.m. by investigators blinded to treatment. Mice were moved from the home vivarium to the behavioral testing room and allowed to acclimate for 1 h before each test.

#### 1) Hindlimb clasping test

Mice were held by the base of the tail and suspended vertically so that all four limbs were clear of surrounding objects. Hindlimb posture was observed for 10 sec and scored on a 0-4 scale ^57^. 4, normal escape extension without clasping; 3, incomplete splay and reduced mobility of one hindlimb with normal toe splay; 2, incomplete splay and reduced mobility of both hindlimbs with normal toe splay; 1, clasping of both hindlimbs with curled, immobile toes; and 0, crossed and immobile forelimbs and hindlimbs with curled toes.

#### 2) Grip strength test

Each mouse was lowered onto the grid until all four paws gripped it. With the torso parallel to the grid, the mouse was pulled steadily backward until it released its grip. Peak grip strength was recorded in three trials and averaged for analysis.

#### 3) Rotarod

Rotarod testing used a six-lane apparatus. Mice were placed on the rod at 4 rpm, and the speed increased to 40 rpm over 5 min. Each mouse completed three trials separated by 10 min, and latency to fall was averaged across trials.

#### 4) Gait analysis

Gait analysis was performed using the VisuGait automated gait-analysis system (BrainTalking, China). Mice were habituated to a glass runway, and forward locomotion was encouraged by food placement at the distal end. Spatial calibration was performed using a standard calibration box (65 × 50 mm), and camera gain and backlight intensity were adjusted to detect paw prints without artifact. Mice were recorded under red-green backlight illumination, and at least three valid consecutive walking trials were acquired per animal (1-4 s/trial). Trials containing fewer than three complete step cycles were excluded. Peak running speed, hindlimb stride length, hindlimb duty cycle, hindlimb swing-phase duration, and diagonal-support duration were extracted using built-in automated analysis and batch-exported for group-level comparison.

### Cell culture and transfection

Human U2OS and SH-SY5Y cells were cultured following the American Type Culture Collection (ATCC) standard guideline. Plasmids were transfected with Lipofectamine 3000 (Invitrogen) according to the manufacturer’s instructions.

### Cellular model construction

U2OS and SH-SY5Y cell models harboring TDP-43 cytosolic condensation were constructed as previously reported^43,44^. Briefly, cells were seeded to 60-70% confluency one day before transfection with mNG-tagged NLS-mutant TDP-43 plasmid (p23-mNG-TDP-43^NLS^). After 24 h, LNP-encapsulated RNA aptamers were added for 6 h, followed by 250 μM sodium arsenite (SA) for 1 h. Cytosolic TDP-43 condensates were imaged by Nikon Spatial Array Confocal microscopy.

### LNP preparation and characterization

Each lipid component, including SM-102, 1,2-distearoyl-sn-glycero-3-phosphocholine (DSPC), cholesterol and 1,2-dimyristoyl-rac-glycero-3-methoxypolyethylene glycol-2000 (DMG-PEG2000) was dissolved in ethanol (Sigma). LNPs were prepared by mixing them with molar ratios of SM-102: DSPC: cholesterol: DMG-PEG2000 at 50:10:38.5:1.5. *In vitro* transcribed RNAs were diluted in pH 4.5 acetate buffer, then encapsulated into LNPs via NanoGenerator Flex-M microfluidic mixing system (PreciGenome) according to manufacturer’s instructions. The LNP-packaged RNAs were then concentrated using Amicon Ultra filters with a molecular weight cutoff of 100 kD (Millipore).

The encapsulation efficiency of RNAs was determined by Quanti-iT RiboGreen RNA assay (Thermo Fisher Scientific) according to manufacturer’s instructions. Dynamic light scattering (Zetasizer Pro, Malvern Panalytical) was used to measure particle size (∼100nm), polydispersity (PDI<0.2) and zeta potential (−10-0 mV).

### Plasmids construction

To construct the plasmids for *in vitro* synthesizing circular RNAs, *Anabaena* PIE carrying T3, T3-M3, poly(A), or other circT3-M3 variant sequences was cloned downstream of the T7 promoter in a pUC57-based plasmid backbone via one-step clone method (TransGen)^19^. All the sequences of RNA templates were shown in Table S1.

To construct recombinant protein purification plasmids, sequence of different TDP-43 variants was cloned into pET9d plasmid via one-step clone method. A TEV cleavage site following the sequence coding MBP tag was introduced downstream of the TDP-43 coding sequence. To construct plasmids expressing HA-tagged TDP-43 variants for co-immunoprecipitation (co-IP) experiments, the HA sequence was cloned downstream of the MBP sequence instead of the His sequence.

To construct the plasmids for expressing TDP-43 in cell, the TDP-43 full-length or the NLS mutated variant was cloned into the p23 plasmid backbone via one-step clone method. The sequence expressing mNG tag was introduced upstream the TDP-43 coding sequence.

All the primer sequences for plasmids construction were listed in Table S2.

### Protein expression and purification

Plasmids expressing different TDP-43-TEV-MBP variants described above were transformed into the *E.coli* expression strain Transetta DE3 (Transgen). Protein expression was induced with 0.5 mM IPTG at 16 °C overnight. Cell pellets were lysed via sonication, and soluble lysates loaded onto MBPTrap HP affinity columns (GE healthcare). After maltose elution, samples were further purified by AKTA size-exclusion chromatography (GE Healthcare) to separate TDP-43 polymers and oligomers and concentrated with Amicon Ultra-4 Centrifugal Filter (Millipore). Purified protein concentrations were determined via modified Bradford assays and verified by SDS-PAGE. Of note, three distinct chromatographic peaks were observed during the purification of TDP-43 protein using size-exclusion chromatography. Based on differences in molecular weight, we separately harvested the highest-molecular-weight peak (left peak) as TDP-43 polymers and the intermediate-molecular-weight peak (middle peak) as TDP-43 oligomers. The purified recombinant proteins were flash frozen with liquid nitrogen and stored at −80°C until use.

### *In vitro* RNA transcription, circularization and purification

The DNA templates for *in vitro* transcription (IVT) were commercially synthesized and received as an insert within the pUC57 plasmid following the T7 promoter (Tsingke). A Xba I restriction site was immediately followed the DNA template sequence.

*In vitro* transcription was conducted using the RiboMax system (Promega). Circular RNA products were treated with *Escherichia coli* poly(A) polymerase (Tinzyme) followed by RNase R (Tinzyme) digestion to degrade linear precursors. RNAs intended for cellular delivery were additionally dephosphorylated with FastAP (Thermo Fisher Scientific) to minimize immunogenicity. All the synthesized RNAs were column purified (NEB), quantified by NanoDrop One spectrophotometer (Thermo Fisher Scientific), and quality-validated on 8 M urea denaturing PAGE.

For making Biotin- and Cy5-labeled RNAs, Biotin-16-UTP (0.25 mM) and Cy5-UTP (0.25 mM) were incorporated into the IVT reaction.

All the primers used for PCR amplification of DNA templates were listed in Table S2.

### RNA isolation and RT-qPCR

Total RNAs from equal number of cultured cells were extracted with Trizol Reagent (Invitrogen) according to the manufacturer’s protocol. The reverse transcription was carried out using PrimeScript RT Master Mix (TaKaRa) according to the manufacturer’s protocol. RT-qPCR was performed using SYBR Green Realtime PCR Master Mix (TOYOBO) and the StepOnePlus real-time PCR system (Applied Biosystems). The relative expression of different sets of genes was quantified to 18s rRNA. All primer sequences for RT-qPCR were listed in Table S2.

### *In vitro* turbidity assay

The RNAs were either denatured or renatured before subjecting to the *in vitro* turbidity assay. To denature the RNAs, appropriate amount of RNAs were diluted in the denaturing buffer (20 mM Tris-HCl pH=7.5, 150 mM NaCl), incubated at 65℃ for 5 min, and immediately placed on ice for at least 2 min. To renature the RNAs, appropriate amount of RNAs were diluted in the renaturing buffer (20mM Tris-HCl pH=7.5, 150 mM NaCl, 10 mM MgCl_2_) or the renaturing buffer with reduced Mg^2+^ (20mM Tris-HCl pH=7.5, 150 mM NaCl, 1 mM MgCl_2_), incubated at 65℃ for 5 min, and slowly cool down (−0.1℃/3 sec) to the RT.

If not otherwise mentioned, for each reaction, 10 µL of 20 µM TDP-43-TEV-MBP protein or its variants was incubated with 10 µL of 2 µM denatured or renatured RNA at RT for 5 min. After that, 1 µL of 72 µM TEV enzyme was added following 15 min incubation at 30℃. Next, 20 µL of the solution was transferred into the 384-well plate (Thermo Fisher Scientific). The turbidity of the solution was detected using the Epoch2 microplate spectrophotometers (BioTek) under the absorbance of OD_600_. The relative turbidity value was calculated by dividing the OD_600_ of the RNA treated sample with the OD_600_ of No RNA treated sample. The RNA that can suppress TDP-43 condensation was determined with the relative turbidity value below 0.5.

### Electrophoretic mobility shift assay (EMSA)

Renatured RNA (2 pmol) was incubated with full-length, mutated or truncated TDP-43 proteins at RNA: protein molar ratios ranging from 1:0 to 1:200 for 15 min at RT. Reactions were resolved on 1.5% agarose or 8% TBE-PAGE gels and visualized via UV transillumination (Tanon).

### Nuclear magnetic resonance (NMR) spectroscopy

NMR experiments were carried out at 298 K on a Bruker 900 MHz spectrometer equipped with a cryogenic probe. Samples were prepared in NMR buffer containing 50 mM PB (pH=7.0), 150 mM NaCl, 2 mM TCEP, and 10% (v/v) D_2_O, with a final volume of 500 μL. ^1^H-^15^N HSQC spectra were recorded for ^15^N-labeled TDP-43 RRM1 upon stepwise addition of T3 RNA to achieve the indicated RNA concentrations. Spectra were acquired using the Bruker standard pulse sequence hsqcetfpf3gpsi with 32 scans. Data were collected with 2048 × 160 complex points in the ^1^H (14 ppm) and ^15^N (21 ppm) dimensions, respectively. All spectra were processed using NMRPipe^58^ and analyzed with Sparky^59^.

### SHAPE probing and SHAPE-MaP reverse transcription

*In vitro* SHAPE probing was performed as described^16,30,31^ with modifications. *In vitro* transcribed and purified RNAs were refolded in 3.3× folding buffer (333 mM HEPES, pH=8.0, 333 mM NaCl, 33 mM MgCl_2_). After refolding, RNAs were incubated with 100 mM of NAI (EMD Millipore) in dimethylsulfoxide (DMSO) for 10 min at 37 °C. For each set of *in vitro* SHAPE probing with proteins, 1.5 pmol of RNAs were incubated with 12 pmol of purified recombinant TDP-43 protein, followed by NAI modification at 100 mM or with DMSO for 10 min at 37 °C. A denaturing control reaction was performed in parallel: RNAs were suspended in a denaturing buffer containing formamide and incubated at 95 °C before modification with NAI. After probing, RNAs were extracted with phenol: chloroform: isoamyl alcohol (25:24:1), ethanol precipitated and dissolved in nuclease-free water.

About 50-200 ng of RNAs were obtained under each treatment and were then reverse transcribed with 200 U of SuperScript II (Invitrogen), 6 mM MnCl_2_ and gene-specific primers for linear or circular RNAs. cDNAs were purified with RNAClean XP (Beckman Coulter). Second-strand synthesis was performed with Q5 hot start high-fidelity DNA polymerase. The resulting PCR products were further isolated with StarPrep Gel Extraction Kit (GenStar). All related primers were listed in Table S2.

### SHAPE-MaP library preparation and sequencing

*In vitro* synthesized linear or circular RNAs at each condition were used as one sample to build a library. In brief, SHAPE-MaP libraries were prepared from 1 ng of DNAs reverse transcribed from SHAPE RNAs, and size-selected with AmpureXP beads (Agencourt) with a 1:1 (bead to sample) ratio to obtain library DNA products spanning 100–400 bp in length. Final libraries were quantified using the Agilent Bioanalyzer 2100. Deep sequencing was performed using the Illumina HiSeq X Ten platform. About 5–15 million sequencing reads were obtained for each sample, with 86% of bases at or above Q30.

### Candidate region screen

Candidate regions were screened from TDP-43 iCLIP datasets (GSM2285561 and GSM2285562). Only exonic binding sites with a false discovery rate (FDR) < 0.05 were retained. A total of 1,646 overlapping sites detected in two replicates were defined as significant sites.

These sites were subsequently merged into regions and ranked by length for further validation according to two criteria: (1) 225 regions were generated by merging sites with intervals of less than 100 nt, and (2) 236 regions were generated by merging sites with intervals of less than 200 nt.

### SHAPE-MaP data analyses

SHAPE-MaP sequencing libraries were prepared by the mixed PCR production. To distinguish reads derived from the same RNA under different treatments, barcodes were added at the 5’ end of reverse primer (Table S2). The barcode and primer were used to separate reads of each RNA in SHAPE-MaP data, by Cutadapt (v.3.10) with the parameter used before^18^.

After read separation, SHAPE-MaP data (NAI-modified, untreated, and denatured control) for each RNA were used to calculate mutation rate (MutR) at single-nucleotide resolution with a minimum efficient coverage of 1,000 (−-min-depth 1,000). SHAPE reactivity was first calculated using the formula [(Modified_MutR_– Untreated_MutR_)/Denatured_MutR_] and then normalized.

For linear RNA, SHAPE-MaP data were directly analyzed by ShapeMapper (V2.1.3) pipeline. For circular RNA, circSHAPE-MaP data were analyzed by CIRCshapemapper (v2) as previously study^19^. In brief, a junction-spanning reference was constructed for circular RNAs, followed by local alignment using Bowtie2. After alignment, the mapped reads would be removed primer sequence, then mutation counts and coverage were determined at single-nucleotide resolution. Finally, SHAPE reactivity was calculated as described above.

### RNA secondary structure modelling

The RNA secondary structures were predicted using RNAfold incorporating SHAPE reactivity values. For linear RNAs, the parameters -p -d2 --shapeMethod=D - shape=SHAPE reactivity < linearRNA.fa were applied. For circular RNAs, the parameters -p -d2 --shapeMethod=D --shape=SHAPE reactivity --circ < circRNA.fa were used. The RNA structures were drawn with StructureEditor.

### *In vitro* protein co-immunoprecipitation (co-IP)

TDP-43-MBP-HA (20 pmol) was incubated with an equal amount of TDP-43-MBP-His (20 pmol) in 200 µL incubation buffer (20 mM Tris-HCl pH 7.5, 150 mM NaCl, 0.05% Tween-20, Roche protease inhibitor cocktail) at 4 °C for 1 h. For RNA-supplemented assays, TDP-43-MBP-HA was first incubated with 2 pmol of renatured RNA aptamers for 1 h before adding TDP-43-MBP-His (20 pmol) for another 1 h. A reverse-order control (RNA added last) was performed in parallel. After incubation, 5% of the mixture was reserved as input, and the remainder was incubated with 20 µL Pierce™ Anti-HA Magnetic Beads (Thermo Fisher) for 2 h at 4 °C, then washed 4 × 5 min with washing buffer (20 mM Tris-HCl pH 7.5, 300 mM NaCl, 0.05% Tween-20). Bound proteins were eluted in 50 µL 1× SDS loading buffer (50 mM Tris pH 6.8, 10% glycerol, 2% SDS, 0.0012% bromophenol blue) at 95 °C for 10 min, and IP/input samples were analyzed by Western blotting.

### Native PAGE WB

The separated oligomers and polymers of full-length, truncated or mutant TDP-43 variants were subjected to Native PAGE experiments. 20 pmol of these purified proteins was individually incubated with 2 pmol renatured RNA aptamers in the incubation buffer (20 mM Tris-HCl pH=7.5, 150 mM NaCl, protease inhibitor cocktail (Roche)) to a total volume of 10 µL at RT for 120 min. After that, the samples were loaded on the NativePAGE^TM^ Bis-Tris Gel (Thermo Fisher Scientific) for electrophoresis at 150V for 60 min and then 250V for 30-60 min at 4℃. Next, the proteins were transferred onto the 0.22 μm PVDF membrane at 250 mA for 80 min, and the membrane was immobilized by 8% acetic acid for 15 min. After that, the membrane was washed with deionized water and underwent air dry. Finally, the membrane was immunoblotted with anti-His followed by ECL chemiluminescence detection. The protein band gray intensity quantification was performed by Image J.

### smTIRF microscopy

All smTIRF data were acquired on a custom-built prism-type TIRF microscope (Olympus IX73) with 532-/637-nm excitation and an EMCCD camera (iXon Ultra 897, Andor), using an OptoSplit II emission splitter (Cairn Research). Laser control was handled by Micro-Manager.

Cy5/biotin-labeled RNA-T3, T9, circT3 or circT3-M3 (1 ng) in 300 µL T50 buffer (20 mM Tris-HCl pH 7.5, 50 mM NaCl, RNase inhibitor) was injected into a custom flow cell (25 µL/min) and immobilized on a streptavidin-coated, PEG-passivated quartz slide; unbound RNA was flushed with imaging buffer A (20 mM Tris-HCl pH 7.5, 0.1 mM DTT, 0.2 mg/ml acetylated BSA (Molecular Cloning Laboratories, BSA-100), 0.0025% P-20 (GE Healthcare, BR100054), 5 mM MgCl2, 100 mM NaCl, RNase inhibitor) supplemented with a photostability/oxygen-scavenging cocktail (saturated Trolox ∼3 mM; PCA–PCD system, 1 mM PCA / 10 nM PCD^60^) to reduce photo-blinking and photobleaching.

For binding activity and dwell time analysis, ∼0.02-1 nM Cy3-TDP-43-MBP in imaging buffer A was injected (100 µL/min) and interactions monitored in real time at ∼23 °C. To measure TDP-43 occupancy per RNA molecule, ∼5-10 nM Cy3-TDP-43-MBP was injected (35 µL/min); after 5 min, imaging buffer B (buffer A without PCA– PCD) was injected (300 µL/min) and Cy3-TDP-43-MBP photobleaching (532 nm, 10 mW) was monitored in real time at ∼23 °C.

### Data analysis of smTIRF imaging

For studies involving Cy5 and biotin labeled RNAs and Cy3-TDP-43-MBP in smTIRF experiments, fluorescent molecules (Cy3-TDP-43-MBP and RNA-Cy5) in two channels were colocalized using a custom written MATLAB script. A 300 ms frame rate with 300 ms laser exposure time was used to examine TDP-43 on RNAs, kymographs were generated along the RNA by a kymograph plug-in in ImageJ. To determine the frequency of TDP-43 on RNAs, single-molecule videos were recorded for 125 sec, Cy3 and Cy5 channels were merged and colocalized molecules with a minimum lifetime of 2 sec were counted as binding events (*N*_binding_). Following real-time single-molecule recording, the number of RNA molecules (*N*_RNA_) was determined by Cy5 dots. The frequencies of RNA binding by TDP-43 (*F*_binding_) were calculated: *F*_binding_ = *N*_binding_ / *N*_RNA_.

Following the real-time single-molecule recording, the binding time of single Cy3-TDP-43-MBP on RNA is defined as τ_on_ and the dissociation time between two binding events on same single RNA is counted as τ_off_. K_on_ and K_off_ is calculated based on τon and τ_off_.

To calculate the Cy3-TDP-43-MBP photobleaching steps on single RNA molecule, single-molecule videos were recorded for 20 min. Photobleaching step analysis was performed using the quickPBSA package or hidden Markov model^61^.

### Biolayer interferometry (BLI)

Biomolecular interactions were measured by biolayer interferometry on a GatorPlus system (GatorBio) with aminopropylsilane (APS) biosensors in accordance with the manufacturer’s protocols. RNA aptamers were immobilized onto APS probes in binding buffer consisting of 1× Dulbecco’s phosphate-buffered saline (DPBS, Gibco). Purified TDP-43 protein acted as the analyte and was diluted to the indicated concentrations in DPBS-T dilution buffer (1× DPBS supplemented with 0.05% (v/v) Tween-20, ABCONE). Assays run at 30°C with shaking at 1,000 rpm and a data acquisition frequency of 10 Hz, following the sequence: baseline–loading–baseline– baseline–baseline–association–dissociation. The first two baseline steps were recorded in DPBS buffer, while the subsequent baseline segments and the dissociation step were conducted in DPBS-T buffer. Raw data are processed with GatorOne software.

### Protein immunofluorescence

U2OS or SH-SY5Y cells were seeded into 8-well plates (Cellvis) for p23-mNG-TDP-43^NLS^ plasmid transfection or TDP-43 cytosolic condensation model construction. After that, the cells were fixed with 4% PFA, permeabilized with Triton X-100, and blocked with 1% BSA. Endogenous G3BP1 was stained with anti-G3BP1 primary antibody (1:200) and Alexa Fluor 555 secondary antibody. Nuclei were counterstained with DAPI. mNG-tagged TDP-43 and immunofluorescent signals were captured by the Nikon Spatial Array Confocal (NSPARC) microscopy (Nikon).

### Amplicon-based circular RNA fluorescence in situ hybridization (abcFISH)

The optimized abcFISH was used to detect the delivered RNA aptamers in cells as described^62^. Briefly, 2×10^4^ U2OS or SH-SY5Y cells were seeded into 18-well plates for transfection/RNA delivery, then fixed with 4% PFA, permeabilized with 0.1% Triton X-100, and equilibrated in equilibration buffer (0.1 mg/mL yeast tRNA, 0.4 U/µL RNasin, 100 mM glycine, 0.1% Tween-20, 2 mM ribonucleoside-vanadyl complexes in 1×PBS). Phosphorylated padlock and primer probes (100 nM each) in hybridization buffer (10% formamide, 2× SSC, 0.1 mg/mL yeast tRNA, 0.2 U/µL RNasin, 20 mM ribonucleoside-vanadyl complexes) were hybridized overnight at 40 °C. Samples were then washed twice with PBSTR (0.1% Tween-20, 0.2 U/µL RNasin in RNase-free 1×PBS) at 37 °C for 20 min each, once with 4× SSC in PBSTR (37 °C, 20 min), and once quickly with PBSTR. Ligation was performed with 0.25 U/µL T4 DNA ligase, 0.5 mg/mL BSA, and 0.4 U/µL RNasin in 1× T4 ligase buffer for 2 h at 25 °C. After washing twice with PBSTR, rolling-circle amplification (RCA) was carried out with 0.5 U/µL Phi29 DNA polymerase, 250 µM dNTPs (TaKaRa, 4019), 0.5 mg/mL BSA, and 0.4 U/µL RNasin in 1× Phi29 buffer at 30 °C for 2 h. Following two PBSTR washes, samples were incubated with fluorescent probes (100 nM each in hybridization buffer) for 1 h at RT, then washed 3 × 10 min with hybridization buffer. G3BP1 and nuclei were stained (DAPI) as above, and samples were mounted in SlowFade Diamond. Images were acquired on a Nikon Spatial Array Confocal (NSPARC) microscope. abcFISH probe sequences are listed in Table S2. Probe sequences used for abcFISH was listed in Table S2.

### RNA-seq analysis

Total RNA-seq reads were trimmed with Trimmomatic^63^ (PE -threads 16 -phred33 TruSeq3PE-2.fa:2:30:10 LEADING:3 TRAILING:3 SLIDINGWINDOW:4:15 MINLEN:30) to remove adapters and low-quality bases, then aligned to rDNA with Bowtie^64^ (v.1.1.2, -m 1 -k 1 -v 2 -p 12 -S); rDNA-mapping reads were discarded. Remaining reads were aligned to hg38 using HISAT2^65^ (v.2.1.0, --known-splicesite-infile --no-softclip --rna-strandness RF --score-min L,-16,0 --mp 7,7 --rfg 0,7 --rdg 0,7 --max-seeds 20 -k 10 -t -p 12 -S) with GENCODE v41 annotation. Gene-level counts were obtained with featureCounts^66^ (−s 2 -p --fraction -O -T 16 -t exon -g gene_name) and converted to FPKM. Differentially expressed genes (DEGs) were identified with the limma^67^ package (|FC| ≥ 2, p < 0.05). Alternative splicing (AS) events were detected using rMATS^68^ (−t paired --readLength 150) from BAM files; significant AS was defined as |ΔPSI| > 0.1 with FDR < 0.05. Differences in AS proportions among groups were assessed by the Kruskal–Wallis test.

### Statistics and reproducibility

Statistics in this study were presented as mean ± SEM for animal behavior experiments (Fig. 6f-n), and mean ± SD for other experiments. Error bars represented SD or SEM in triplicate experiments if not mentioned otherwise. Statistical significance for comparisons was generally assessed by Student’s t test. *p-values* below 0.05 were marked by one asterisk, while two asterisks indicate *p-value* < 0.01, three asterisks indicate *p-value* < 0.001, and four asterisks indicate *p-value* <0.0001.

## DATA AND MATERIALS AVAILABILITY

All data are available in the main text or the supplementary materials. This paper analyzes existing, publicly available data (GSM2285561 and GSM2285562) deposited at the Gene Expression Omnibus. All the materials generated in this study are available from the lead contact with a completed Materials Transfer Agreement.

## ACKNOWLEDGMENTS

We thank all members of the Chen lab for their comments and suggestions throughout this project. We thank Prof. Li Yang and Dr. Pei-Hong Zhang for helping with RNA sequencing and analysis. We thank Zhijun Liu from the Nuclear Magnetic Resonance System (https://cstr.cn/31129.02.NFPS.NMRSystem) at the National Facility for Protein Science in Shanghai (https://cstr.cn/31129.02.NFPS) for providing technical support and assistance in NMR data collection and analysis. This work is supported by the Strategic Priority Research Program of the CAS (XDB0570000), the National Key R&D Program of China (2021YFA1300500 and 2021YFA1300501, 2024YFC3405900 and 2024YFC3405902, 2025YFA1308800), the Science and Technology Commission of Shanghai Municipality (23DX1900100 and 23DX1900101), the National Natural Science Foundation of China (32571501, 32301079, 22425704, 82188101). This work has been supported by the New Cornerstone Science Foundation through the New Cornerstone Investigator Program and the Shanghai Municipal Science and Technology Major Project. L.-L.C. is a SANS senior investigator. H.W. acknowledges the support from Youth Innovation Promotion Association, CAS.

## AUTHOR CONTRIBUTIONS

L.-L.C. conceived the project; L.-L.C. and H.W. designed experiments; H.W., P.-F.L., L.L. and J.-W.C. performed all the molecular, biochemical, and cellular experiments with the help of Y.-Y.P, X.W., J.L., Y.-X.L. and G.X, supervised by L.-L.C. and H.W.; J.H. and Y. W. performed the NMR experiments supervised by C.L.; H.W., B.-W.J., J.H. and Y.W. performed the animal experiments supervised by C.L.; F.N. performed computational analyses supervised by L.-L.C; L.-L.C. and H.W. drafted the manuscript, L.-L.C., H.W. and C.L. edited the manuscript.

## SUPPLEMENTARY FIGURES AND FIGURE LEGENDS

**Fig. S1.**
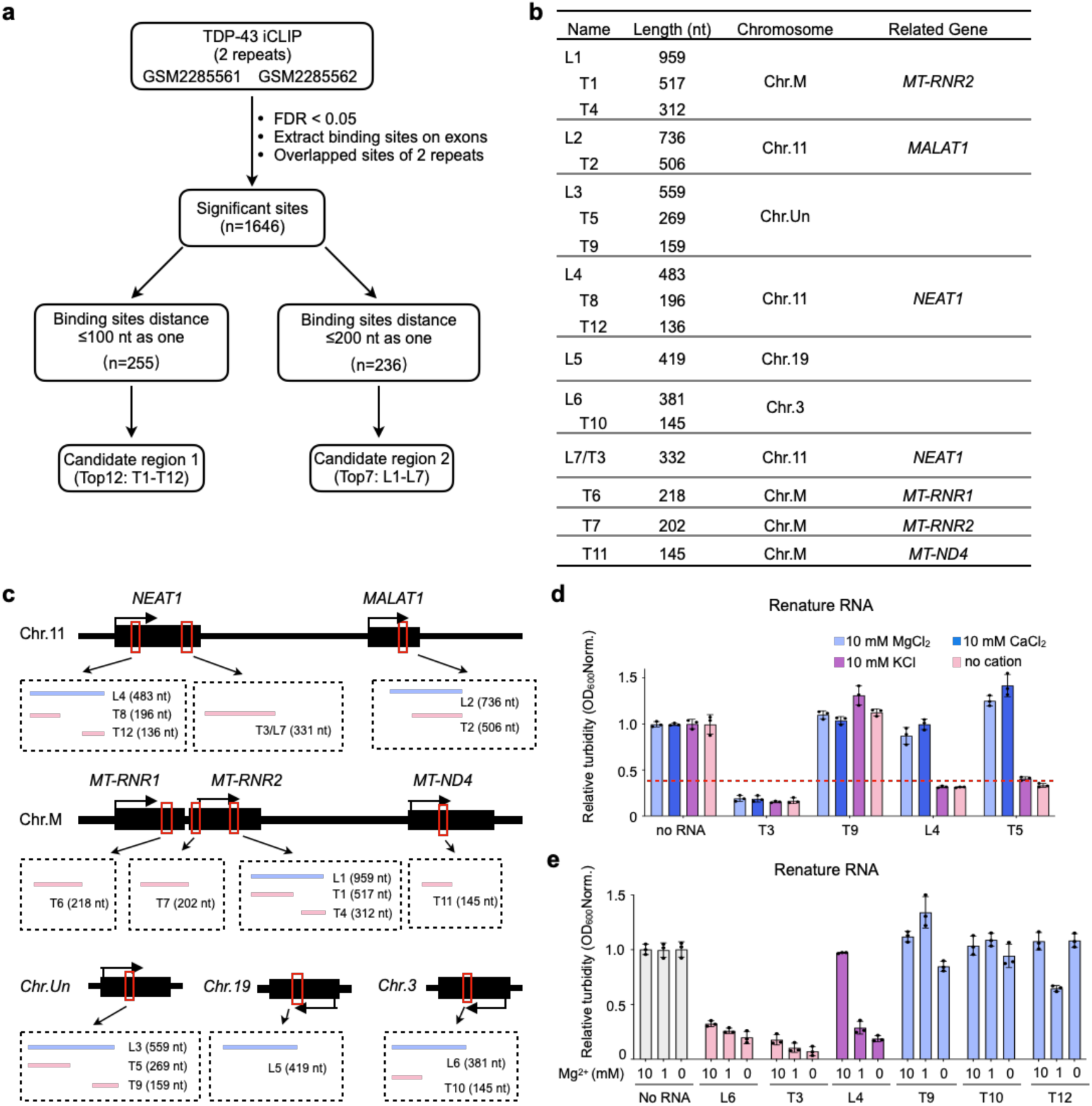
Denatured TDP-43-interacting RNAs inhibit TDP-43 condensation *in vitro.* **a** Schematic for identifying TDP-43 interacting RNA fragments from the TDP-43 iCLIP-seq data (GSM2285561 and GSM2285562)^24^. **b** The length, chromosome location, and related genes of the TDP-43 interacting RNA fragments identified by analyzing the TDP-43 iCLIP-seq data with the pipeline of S1A. **c** Genomic localization of the 18 identified TDP-43 interacting RNAs. **d** 10 mM of divalent cation (Mg^2+^ or Ca^2+^) is sufficient to block the efficacy of L4 and T5 RNAs, but not T3 RNA for mitigating TDP-43 condensation. T9 RNA that shows no efficacy in both denature and renature conditions is also tested. 10 mM of K^+^ or no cation condition is used as negative controls. Oligomeric MBP-tagged TDP-43 and renatured RNAs are used in this experiment. **e** 1 mM Mg^2+^ which mimics the concentration in cell does not affect L4 RNA’ capacity for preventing TDP-43 condensation. L6 and T3 RNAs’ efficacy is not affected by the ion concentration. T9, T10, and T12 RNAs have no efficacy to suppress TDP-43 condensation at different concentration of ion. Oligomeric MBP-tagged TDP-43 and renatured RNAs are used in this experiment.

**Fig. S2.**
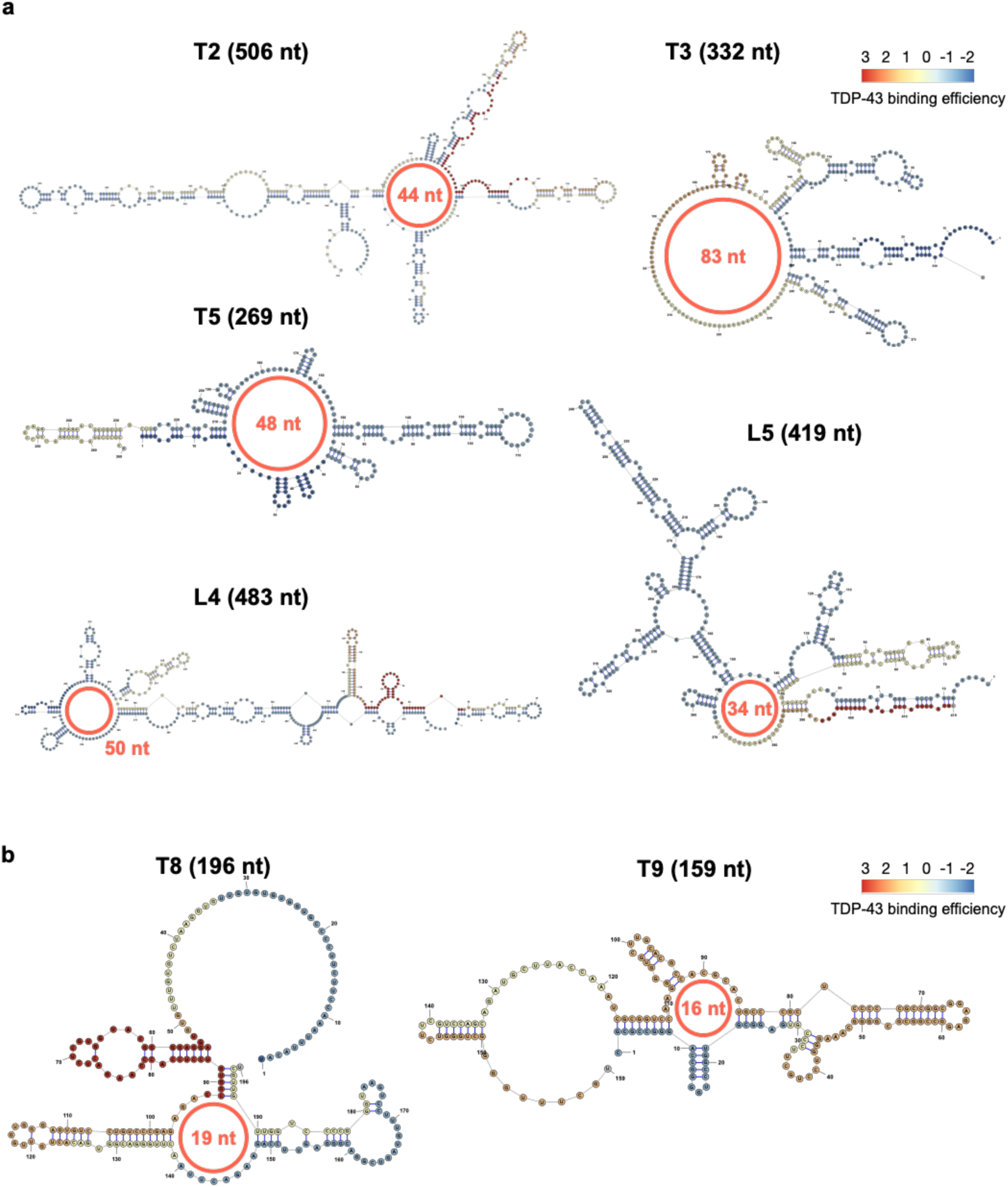
Active TDP-43-interacting RNAs contain large multi-branched loops. a, **b** RNA secondary structures of T2, T3, T5, L4, and L5 that show efficacy in suppressing TDP-43 condensation (**a**) and T8, T9 that do not have this activity (**b**). All structures are modeled with SHAPE-MaP experiments. The multi-branched loops are marked in red.

**Fig. S3.**
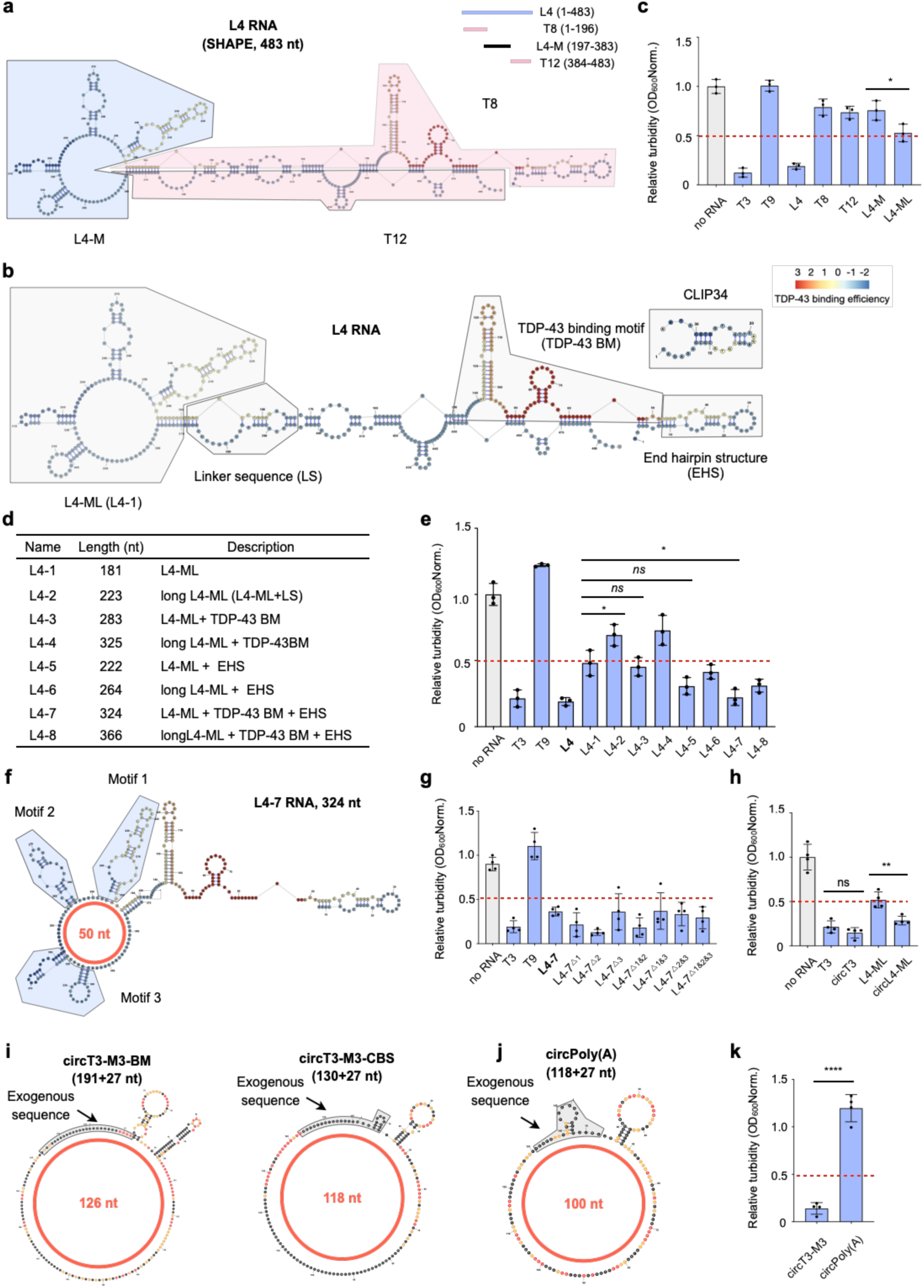
The loosen structure is essential for the anti-TDP-43 condensation function. **a** Secondary structure of L4 RNA modeled with SHAPE-MaP experiment. L4 RNA is architecturally organized by T8 and T12 (pink) flanked with the L4-M domain with a 50-nt multi-branched loop (blue). **b** L4 RNA contains a loosely L4-ML structure, a liner sequence (LS), a TDP-43 binding motif (TDP-43 BM), and an end hairpin structure that is similar to *Clip34*. These substructures are marked in grey. **c** Turbidity assay shows that the L4-ML but not the L4-M has the capacity to suppress TDP-43 condensation. T3 and L4 RNAs are used as positive controls; T8, T9, and T12 RNAs are used as negative controls. Oligomeric MBP-tagged TDP-43 and renatured RNAs are used in this experiment. **d** The architecture of L4 RNA truncations. **e** Turbidity assay to detect the efficacy of L4 RNA truncations. T3 and T9 RNA are used as positive and negative controls, respectively. L4-7 shows the best efficacy among all the truncations. Oligomeric MBP-tagged TDP-43 and renatured RNAs are used in this experiment. **f** The predicted structure of L4-7 RNA, which is artificialized by combining the L4-ML, the TDP-43 binding sequence (BM), and the end hairpin structure (EHS). Three hairpins in the non-structured multi-branched loop are marked in blue. **g** Individual or combined deletion of the hairpin structures do not affect the efficacy of L4-7, revealed by the turbidity assay. T3 RNA is used as a positive control, T9 RNA is used as a negative control. Oligomeric MBP-tagged TDP-43 and renatured RNAs are used in this experiment. **h** The circularized T3 (circT3) has comparable efficacy to linear T3, while circularized L4-ML (circL4-ML) has better efficacy than linear L4-ML in suppressing TDP-43 condensation, revealed by the turbidity assay. Oligomeric MBP-tagged TDP-43 and renatured RNAs are used in this experiment. **i, j** Secondary structures of circT3-M3-BM and circT3-M3-CBS (**i**), and circPoly(A) (**j**) modeled with SHAPE-MaP experiments. circT3-M3-BM contains a 126-nt multi-branched loop structure, circT3-M3-CBS contains a 118-nt multi-branched loop structure, and circPoly(A) contains a 100-nt multi-branched loop structure. The exogenous 27 nt sequence generated during circularizing process is marked in grey. **k** circPoly(A) shows no efficacy in suppressing TDP-43 condensation, revealed by the turbidity assay. circT3-M3 is used as a positive control. Oligomeric MBP-tagged TDP-43 and renatured RNAs are used in this experiment.

**Fig. S4.**
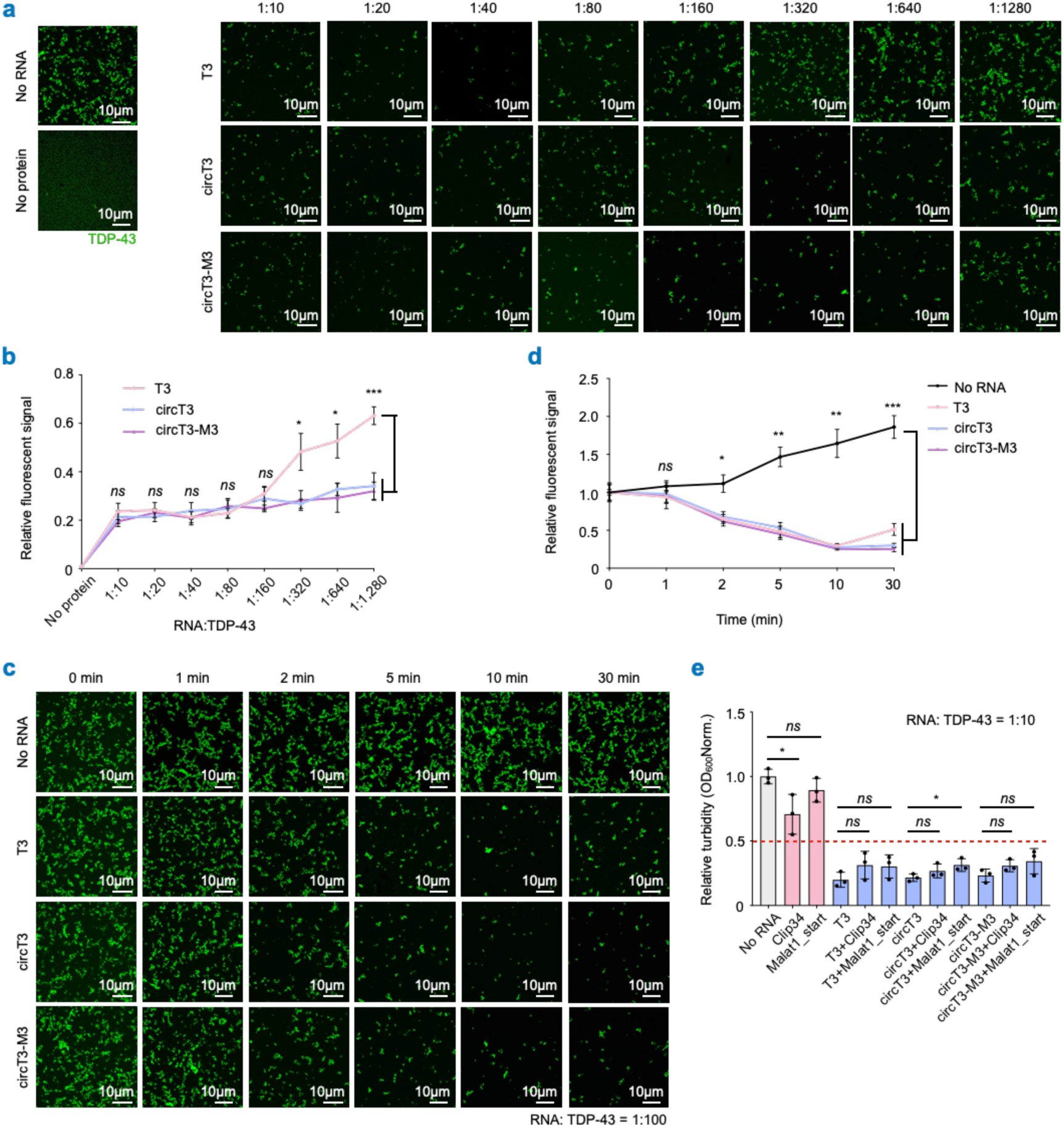
Circular RNA aptamers suppress TDP-43 condensation sub-stoichiometrically and in a time-dependent manner. **a** circT3 and circT3-M3 suppress TDP-43-EGFP condensation sub-stoichiometrically and outcompete linear T3 at low dosage, revealed by the imaging assays. **b** Statistical analysis of the condensed TDP-43-EGFP signal from (**a**), normalized to no RNA-treated group. **c** At the RNA: TDP-43 ratio of 1:100, T3, RNA aptamers start to exhibit efficacy in suppressing TDP-43-EGFP condensation at 2 min of incubation, revealed by the imaging assays. **d** Statistical analysis of the condensed TDP-43-EGFP signal from (**c**). **e** Turbidity assay reveals that *Clip34* exerts weak efficacy in suppressing TDP-43 condensation, whereas *Malat1_start* was ineffective at the RNA: TDP-43 ratio of 1:10. Combining either RNA with T3/circT3/circT3-M3 fails to synergistically improve the anti-TDP-43 condensation efficacy.

**Fig. S5.**
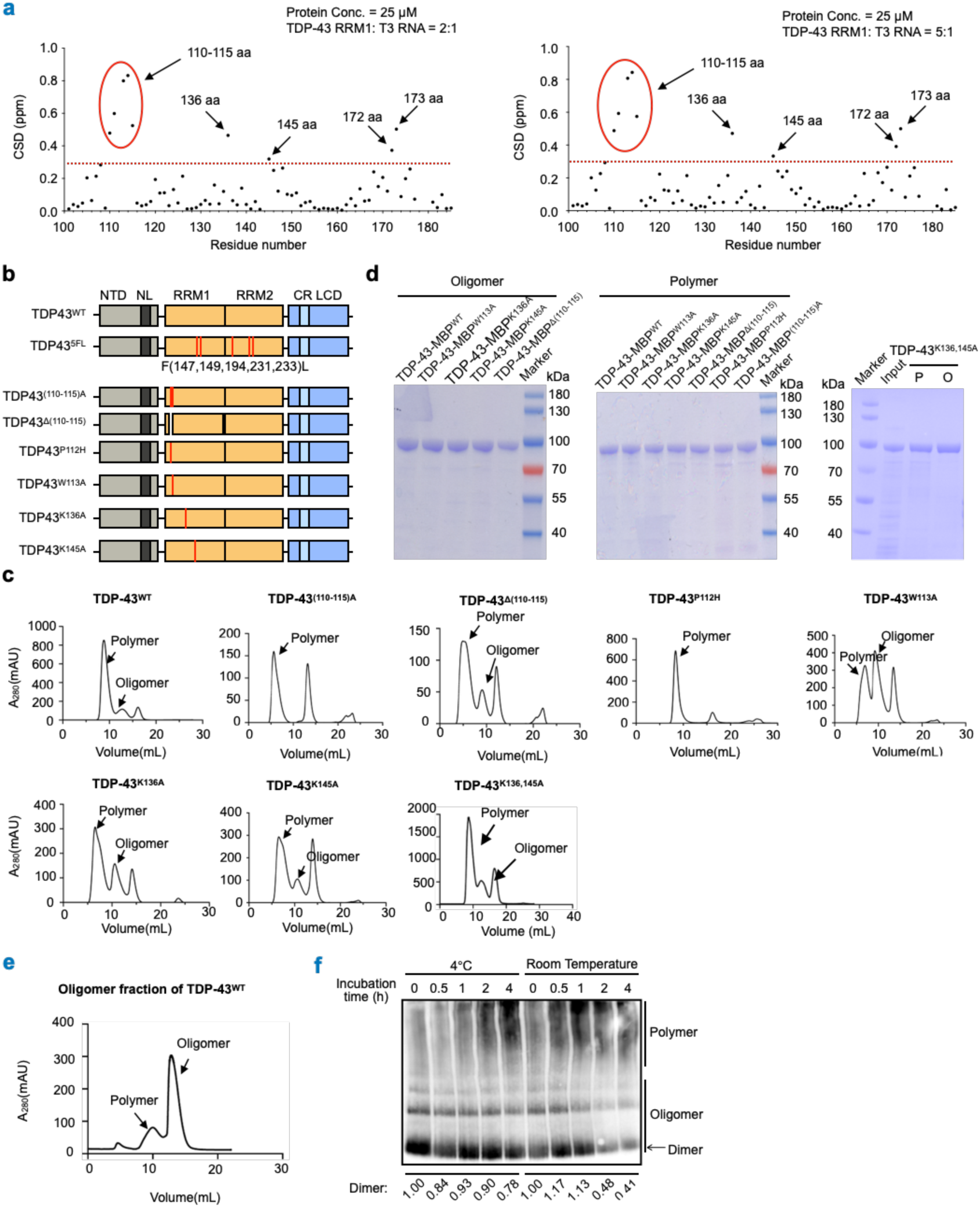
Construction and purification of TDP-43 mutation oligomers and polymers. **a** NMR chemical shift deviations (CSDs) in TDP-43 RRM1 after addition of T3 at protein: RNA ratios of 2:1 (left) and 5:1 (right). Residues with CSDs above 0.3 ppm are highlighted as candidate interaction sites. **b** Domain architecture of TDP-43 mutations. The mutated RNA binding sites are marked in red. **c** AKTA purification of MBP-tagged TDP-43 wild-type, (110-115)A, △(110-115), P112H, W113A, K136A, K145A, and K136,145A variants. The indicated oligomers and polymers are separated. **d** Bradford staining of the purified MBP-tagged TDP-43 variants. The oligomers and polymers are separated. **e** Oligomeric state verification of wild-type TDP-43-MBP oligomers obtained via AKTA purification. **f** Purified TDP-43 oligomers gradually convert into polymers in a time-dependent manner, revealed by native-PAGE.

**Fig. S6.**
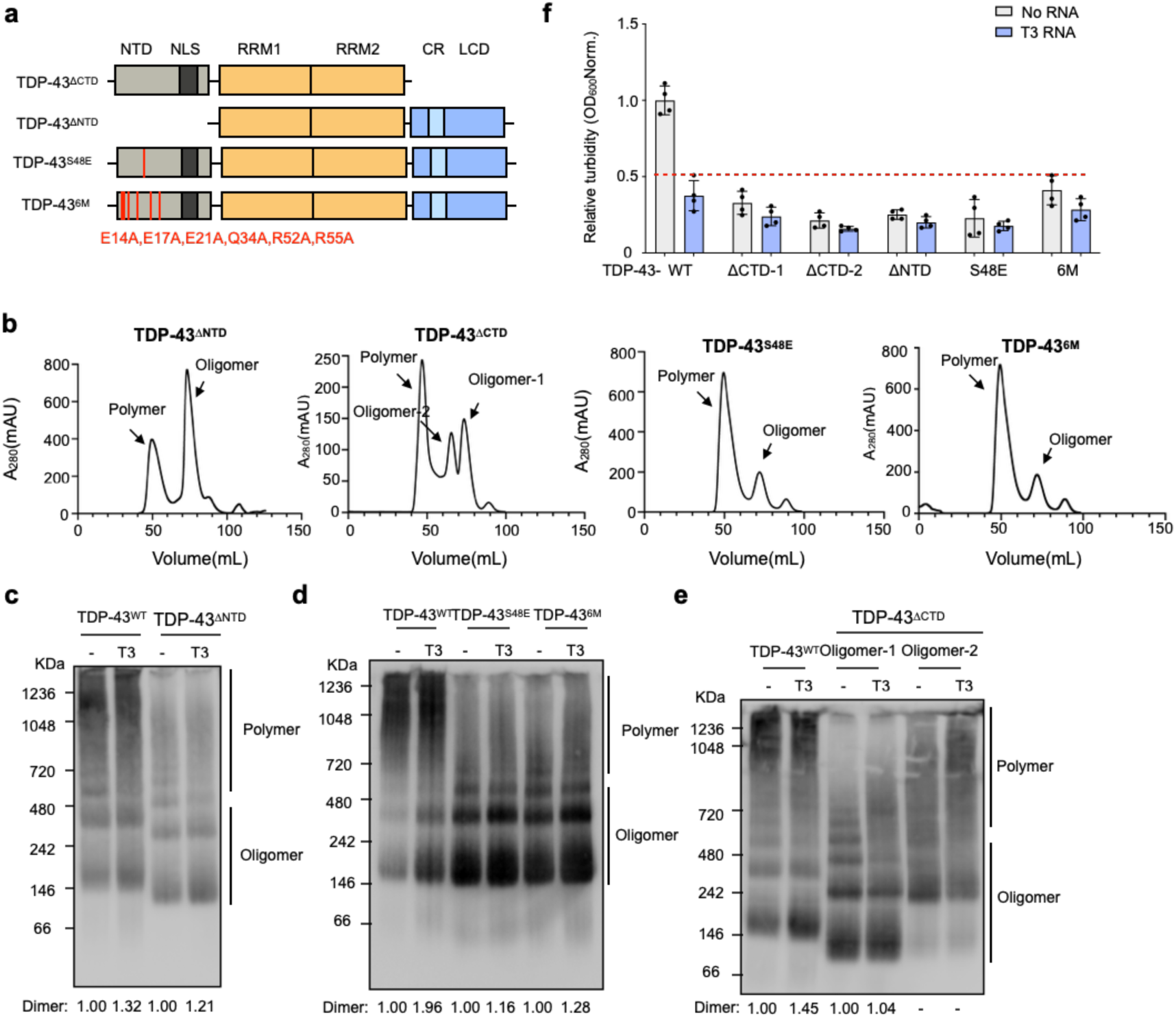
TDP-43 N-terminal and C-terminal mutations or truncations form oligomers that do not condensate. **a** Domain architecture of TDP-43 C-terminal and N-terminal deleted truncations, as well as S48E and 6M mutations. The mutated sites are marked in red. **b** AKTA purification of MBP-tagged TDP-43 △NTD, △CTD truncations, S48E and 6M mutations. The indicated oligomers and polymers are separated. **c-e** Native PAGE of MBP-tagged TDP-43^△NTD^ (**c**), TDP-43^S48E^ and TDP^6M^ (**d**), and TDP-43^△CTD^ (**e**) oligomers maintained at RT for 2h. All of them tend to stay at oligomeric state. MBP-tagged wild-type TDP-43 oligomer is used as control, which is converted into polymers after 2 h incubation. T3 RNA inhibits the polymerization of wild-type TDP-43, but has no impact to these TDP-43 truncations and mutations. **f** MBP-tagged TDP-43^△NTD^, TDP-43^△CTD^, TDP-43^S48E^ and TDP^6M^ oligomers are soluble, regardless of the addition of T3 RNA aptamer, revealed by the turbidity assay. MBP-tagged wild-type TDP-43 oligomer is used as a control.

**Fig. S7.**
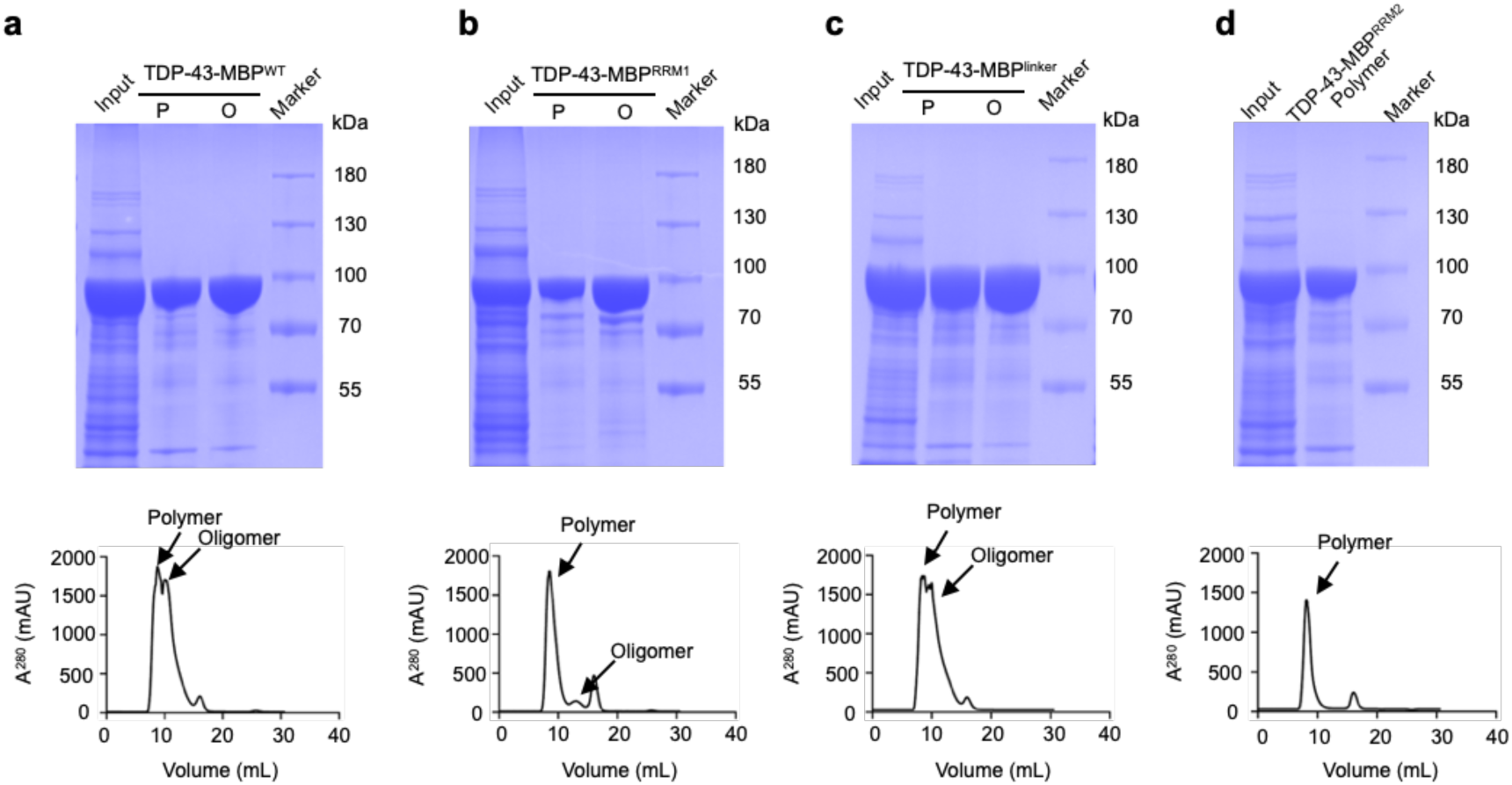
Purification of TDP-43 RRM1, linker, and RRM2 mutants. **a-d** Purification of MBP-tagged TDP-43^WT^ (**a**), TDP-43^RRM1^ (**b**), TDP-43^liner^ (**c**), and TDP-43^RRM2^ (**d**). The oligomers and polymers are separated. (Upper) The Bradford staining of the purified proteins. P: polymers, O: oligomers. (Bottom) The AKTA purification of these TDP-43 variants.

**Fig. S8.**
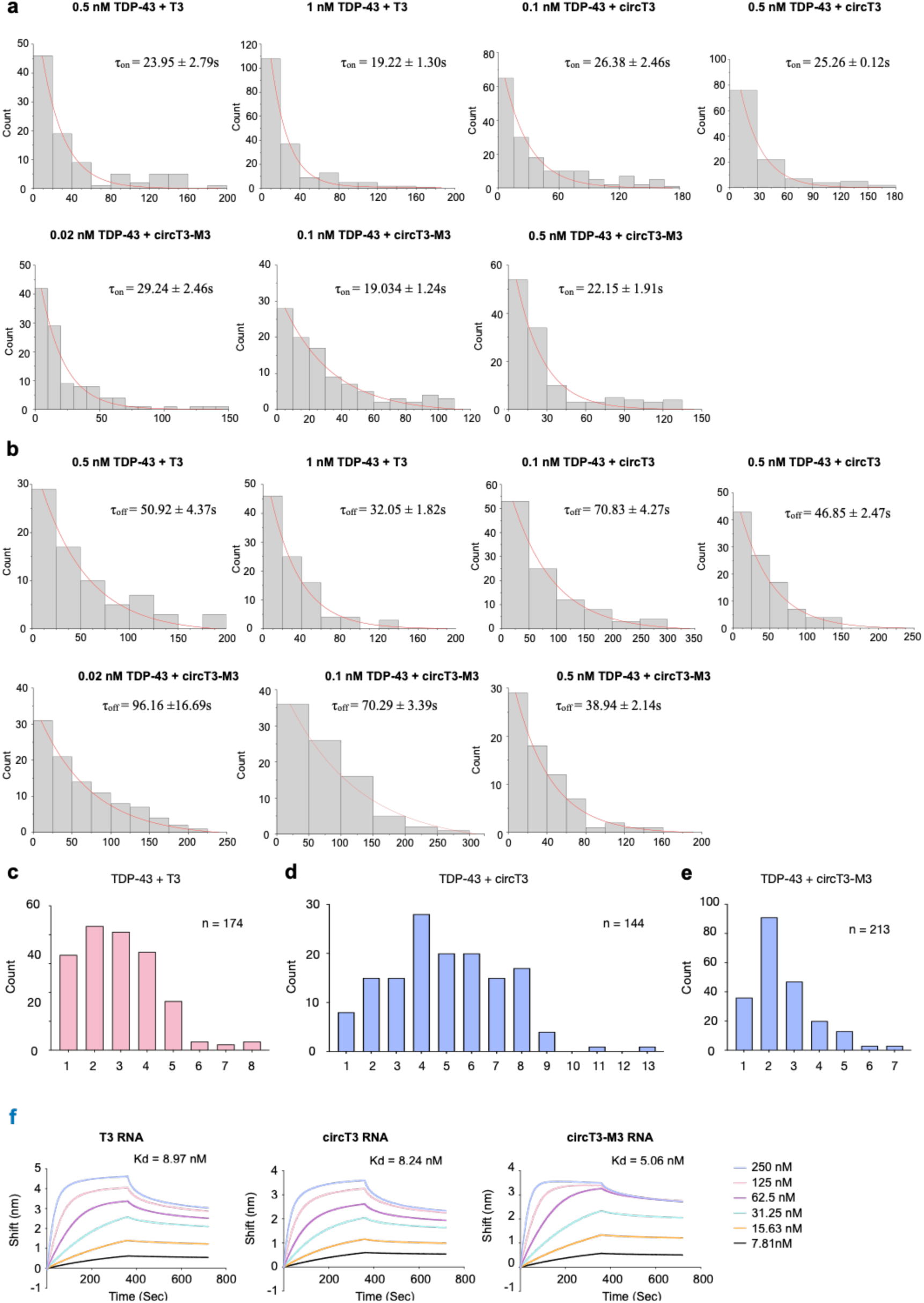
RNA aptamers transiently bind multiple TDP-43 molecules. **a** The association time (τ_on_) calculation between MBP-tagged TDP-43 and RNA aptamers. 0.5 nM and 1 nM TDP-43-MBP are used for detecting the interaction with T3 and circT3; 0.02 nM, 0.1 nM, and 0.5 nM TDP-43-MBP are used for detecting the interaction with circT3-M3. **b** The dissociation time (τ_off_) calculation between MBP-tagged TDP-43 and RNA aptamers. 0.5 nM and 1 nM TDP-43-MBP are used for detecting the interaction with T3 and circT3; 0.02 nM, 0.1 nM, and 0.5 nM TDP-43-MBP are used for detecting the interaction with circT3-M3. **c-e** The number of TDP-43 bindings on a single T3 (**c**), circT3 (**d**), and circT3-M3 (**e**) molecule is calculated based on the step analysis of photobleaching from single-molecule fluorescence imaging of TDP-43-MBP-Cy3. n, number of events examined. **f** Apparent binding affinity estimation between RNA aptamers and TDP-43-MBP protein via biolayer interferometry. The titrated TDP-43 concentrations varying from 250 nM to 7.81 nM are evaluated.

**Fig. S9.**
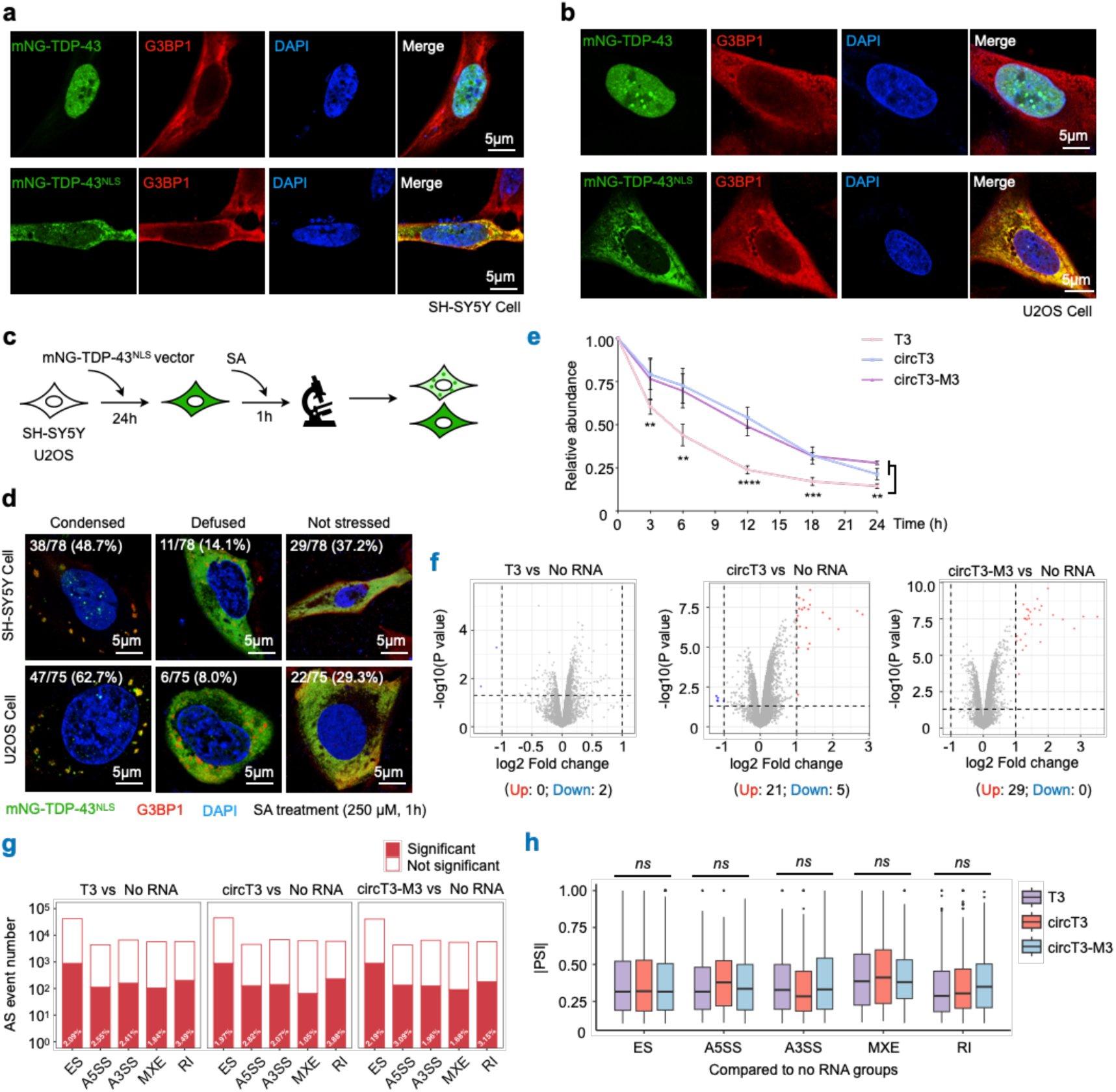
Construction of cellular models exhibiting cytosolic TDP-43 condensation. **a, b** The localization of ectopically expressed mNG-TDP43 and mNG-TDP-43^NLS^ mutation in SH-SY5Y (**a**) and U2OS (**b**) cells. **c** Schematic of constructing the SH-SY5Y and U2OS cell models that contain cytosolic TDP-43 condensation. **d** Representative images showing the SH-SY5Y and U2OS cell models constructed with the pipeline from **c**. The percentages of cells displaying condensed or diffused TDP-43 signals, along with unstressed cells are quantified. mNG-TDP-43 (green), G3BP1 (red). The nucleus is stained with DAPI. Scale bar = 5 μm. **e** Intracellular stability of RNA aptamers in wild-type SH-SY5Y cells, revealed by RT-qPCR. The expression levels of the tested RNAs are normalized to the expression of *GAPDH*. **f** Dysregulated genes in RNA aptamer-treated SH-SY5Y cells. The upregulated genes are marked in red and downregulated genes are marked in blue. Cutoff: *p-value*<0.05, fold change>2 or <0.5. **g** Very limited alterations in the number of alternative splicing (AS) events in RNA aptamer-treated SH-SY5Y cells. Cutoff: False Discovery Rate (FDR)<0.05, |percent spliced in (PSI)|>0.1. **h** Proportion of significantly changed AS events does not differ among comparisons (Kruskal–Wallis test).

**Fig. S10.**
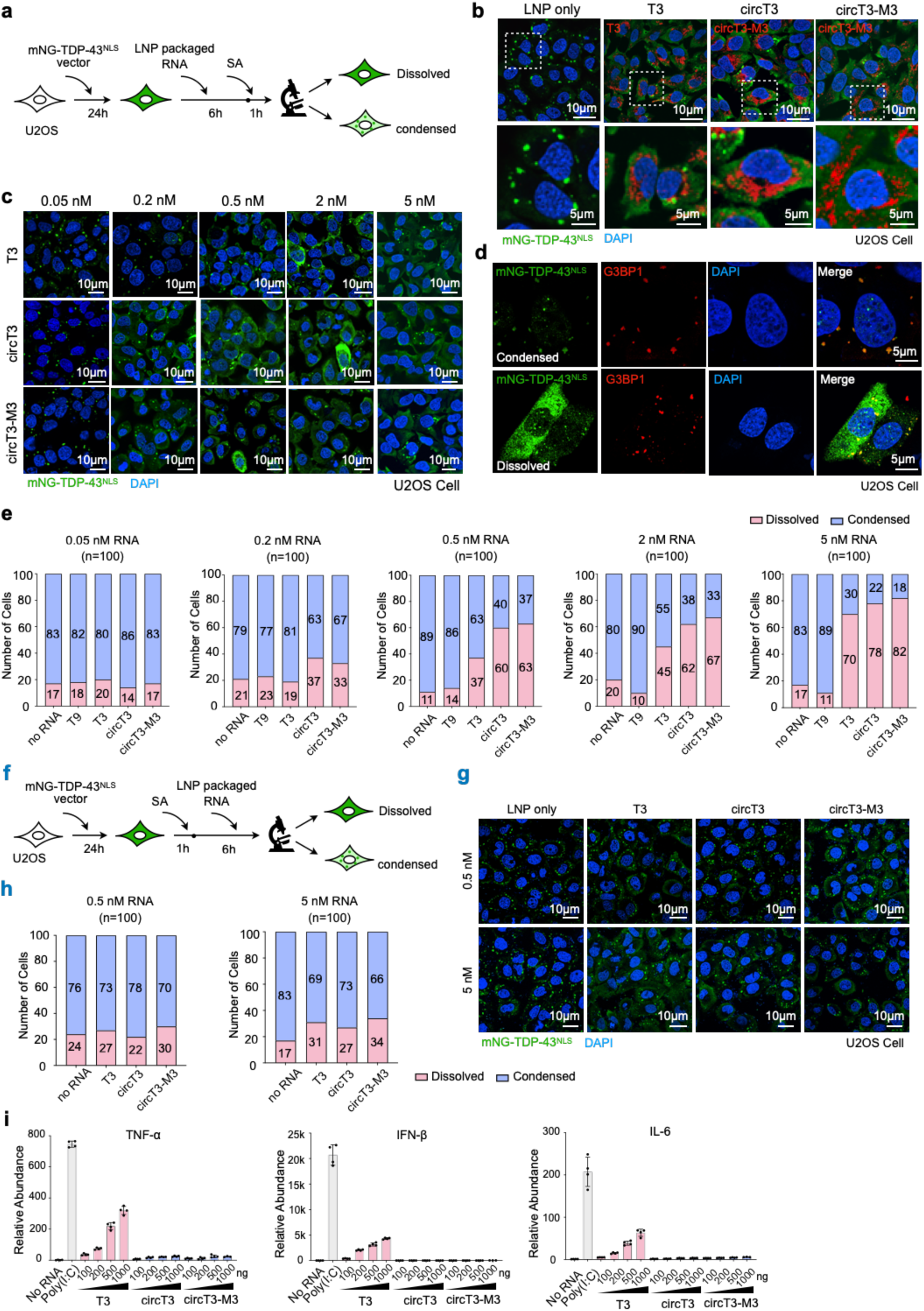
Circular RNA aptamers suppress TDP-43 condensation in U2OS cells. **a** Schematic of employing LNP-packaged RNA aptamers to suppress TDP-43 cytosolic condensation in the U2OS cell model. **b** Representative images showing the delivered T3, circT3, and circT3 RNAs in the U2OS cell model. mNG-TDP-43 (green), RNA aptamers (red). The nucleus is stained with DAPI. Scale bar = 5 μm. **c** Representative images showing the condensation state of TDP-43 post RNA aptamer delivery at concentrations of 0.05 nM, 0.2 nM, 0.5 nM, 2 nM, and 5 nM in the U2OS cell model. mNG-TDP-43 is stained in green. The nucleus is stained with DAPI. Scale bar = 10 μm. **d** Representative zoom-in images showing the condensed (upper) and dissolved (bottom) TDP-43 post RNA aptamer delivery in the U2OS cell model. mNG-TDP-43 (green), G3BP1 (red). The nucleus is stained with DAPI. Scale bar = 5 μm. **e** Statistics from (**c**) showing that Capts exhibit inhibitory efficacy to TDP-43 condensation start from the concentration of 0.2 nM, outperforming T3 which initiates efficacy that start from 0.5 nM. **f** Schematic of employing LNP-packaged RNA aptamers to reverse the pre-formed TDP-43 condensates in U2OS cells. **g** Representative images showing the condensation state of TDP-43 post RNA aptamer delivery at 0.5 nM and 5 nM. mNG-TDP-43 is stained in green. The nucleus is stained with DAPI. Scale bar = 10 μm. **h** Statistics from (**g**) showing that the RNA aptamers exhibit limited capacity to reverse pre-formed TDP-43 condensates at 0.5 nM and 5 nM concentrations. **i** T3 RNA delivery elicits a moderate but reduced inflammatory response compared to poly(I:C) stimulation, as indicated by elevated TNF-α, IFN-β, and IL-6. While circT3 and circT3-M3 delivery induces minimal cytokine expression.

**Fig. S11.**
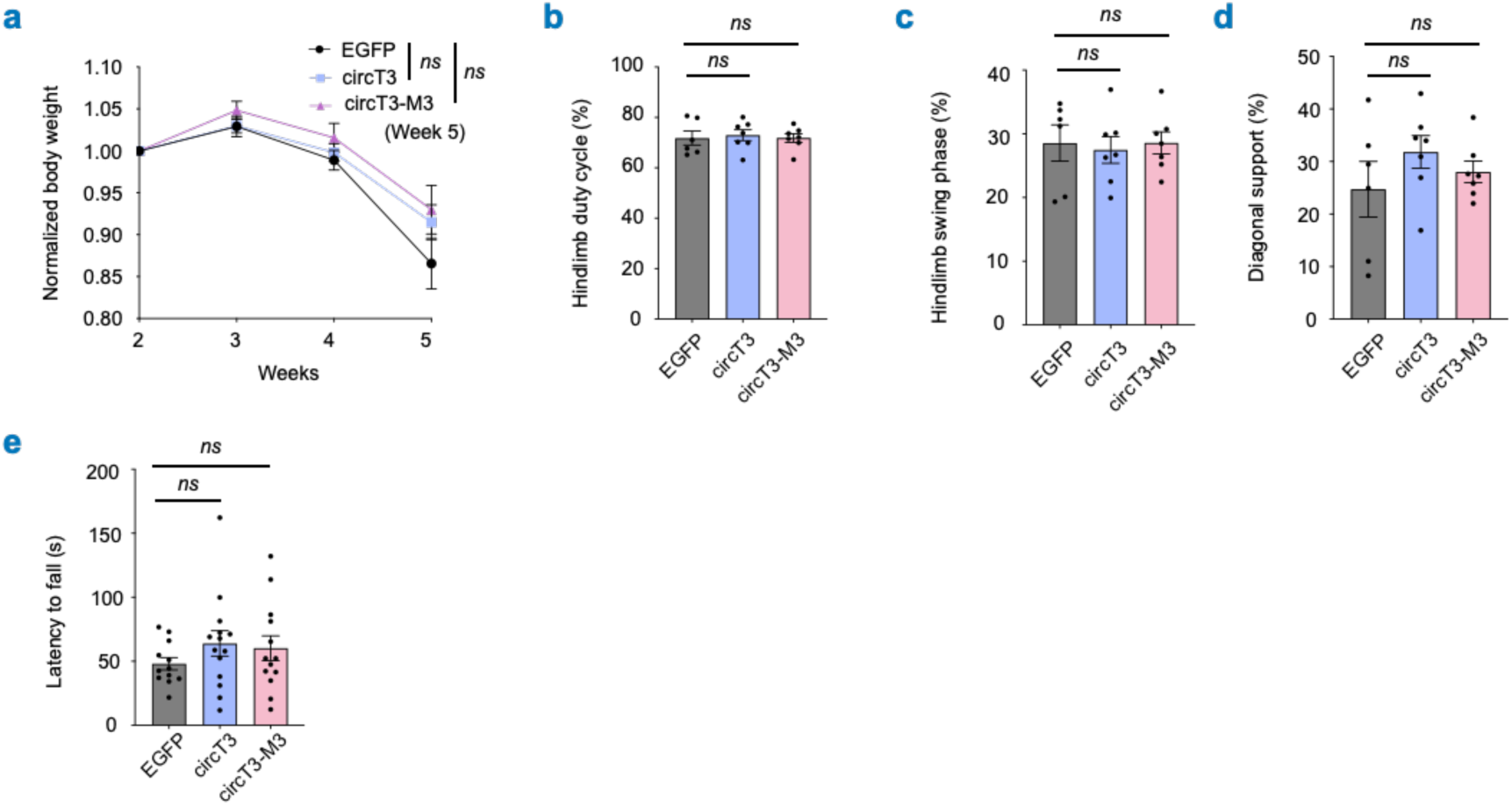
Circular RNA aptamers fail to rescue higher-order behavioral performance in the mouse model. **a** Relative body weight of the AAV-treated NEFH-tTA/tetO-hTDP-43^ΔNLS^ mice, normalized to the averaged body weight of the EGFP-expressing group at week 2. **b-d** Gait analysis reveals that Capt-treated mice show hindlimb duty cycle (**b**), hindlimb swing-phase duration (**c**), and diagonal-support duration (**d**) comparable to those of EGFP-expressing mice. **e** Rotarod assays reveal that the latency-to-fall is not rescued in Capt-treated mice.

**Fig. S12.**
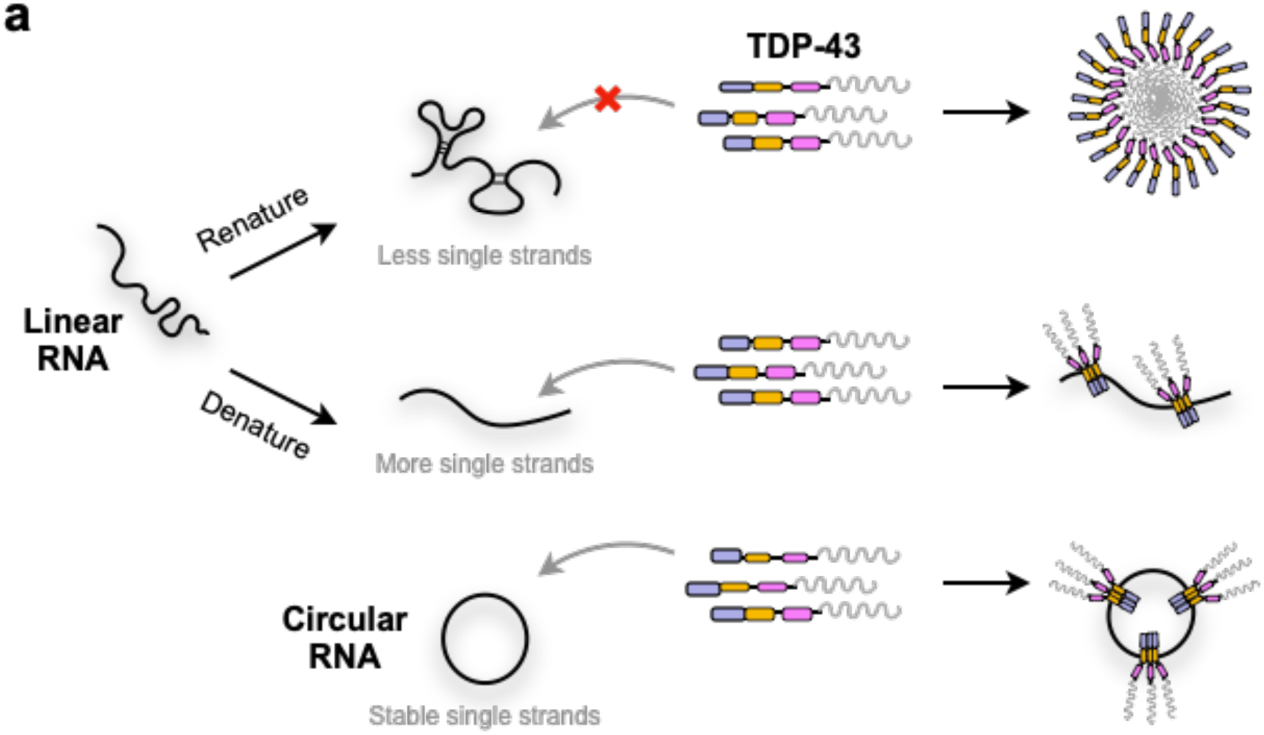
RNA structures facilitate the effective inhibition of TDP-43 condensation. **a** Schematic to show that the denatured linear RNAs offer more single-stranded regions than their renatured formats to engage more TDP-43 molecules, thereby preventing the C-terminal-mediated condensation. Circular RNAs with stable single-stranded regions have the highest capacity to inhibit TDP-43 condensation.

## REFERENCES

1 Ross, C. A. & Poirier, M. A. Protein aggregation and neurodegenerative disease. Nat Med 10 Suppl, S10–17 (2004).

2 Valastyan, J. S. & Lindquist, S. Mechanisms of protein-folding diseases at a glance. Dis Model Mech 7, 9–14 (2014).

3 Maharana, S. et al. RNA buffers the phase separation behavior of prion-like RNA binding proteins. Science 360, 918–921 (2018).

4 Neumann, M. et al. Ubiquitinated TDP-43 in frontotemporal lobar degeneration and amyotrophic lateral sclerosis. Science 314, 130–133 (2006).

5 Ling, S. C., Polymenidou, M. & Cleveland, D. W. Converging mechanisms in ALS and FTD: disrupted RNA and protein homeostasis. Neuron 79, 416–438 (2013).

6 Yan, X. et al. Intra-condensate demixing of TDP-43 inside stress granules generates pathological aggregates. Cell 188, 4123–4140 e4118 (2025).

7 Babazadeh, A., Rayner, S. L., Lee, A. & Chung, R. S. TDP-43 as a therapeutic target in neurodegenerative diseases: Focusing on motor neuron disease and frontotemporal dementia. Ageing Res Rev 92, 102085 (2023).

8 Hayes, L. R. & Kalab, P. Emerging Therapies and Novel Targets for TDP-43 Proteinopathy in ALS/FTD. Neurotherapeutics 19, 1061–1084 (2022).

9 Afroz, T. et al. Functional and dynamic polymerization of the ALS-linked protein TDP-43 antagonizes its pathologic aggregation. Nat Commun 8, 45 (2017).

10 Mercado, P. A., Ayala, Y. M., Romano, M., Buratti, E. & Baralle, F. E. Depletion of TDP 43 overrides the need for exonic and intronic splicing enhancers in the human apoA-II gene. Nucleic Acids Res 33, 6000–6010 (2005).

11 Hruska-Plochan, M. et al. A model of human neural networks reveals NPTX2 pathology in ALS and FTLD. Nature 626, 1073–1083 (2024).

12 Zhou, J. & Rossi, J. Aptamers as targeted therapeutics: current potential and challenges. Nat Rev Drug Discov 16, 181–202 (2017).

13 Copley, K. E. et al. Short RNA chaperones promote aggregation-resistant TDP-43 conformers to mitigate neurodegeneration. Science 392, eadv3301 (2026).

14 Enuka, Y. et al. Circular RNAs are long-lived and display only minimal early alterations in response to a growth factor. Nucleic Acids Res 44, 1370–1383 (2016).

15 Zhang, Y. et al. The Biogenesis of Nascent Circular RNAs. Cell Rep 15, 611–624 (2016).

16 Liu, C. X. et al. Structure and Degradation of Circular RNAs Regulate PKR Activation in Innate Immunity. Cell 177, 865–880 e821 (2019).

17 Liu, C. X., Yang, L. & Chen, L. L. Dynamic conformation: Marching toward circular RNA function and application. Mol Cell 84, 3596–3609 (2024).

18 Liu, C. X. et al. RNA circles with minimized immunogenicity as potent PKR inhibitors. Mol Cell 82, 420–434 e426 (2022).

19 Guo, S. K. et al. Therapeutic application of circular RNA aptamers in a mouse model of psoriasis. Nat Biotechnol 43, 236–246 (2025).

20 Guo, S. K. et al. Therapeutic circRNA aptamer alleviates PKR-associated osteoarthritis. Sci Bull (Beijing) 70, 2232–2236 (2025).

21 Feng, X. et al. Circular RNA aptamers targeting neuroinflammation ameliorate Alzheimer disease phenotypes in mouse models. Nat Biotechnol 44, 454–463 (2026). 10.1038/s41587-025-02624-w

22 Rummens, J. et al. TDP-43 seeding induces cytoplasmic aggregation heterogeneity and nuclear loss of function of TDP-43. Neuron 113, 1597–1613 e1598 (2025).

23 Wu, M. et al. lncRNA SLERT controls phase separation of FC/DFCs to facilitate Pol I transcription. Science 373, 547–555 (2021).

24 Wu, H. et al. Unusual Processing Generates SPA LncRNAs that Sequester Multiple RNA Binding Proteins. Mol Cell 64, 534–548 (2016).

25 Polymenidou, M. et al. Long pre-mRNA depletion and RNA missplicing contribute to neuronal vulnerability from loss of TDP-43. Nat Neurosci 14, 459–468 (2011).

26 Tollervey, J. R. et al. Characterizing the RNA targets and position-dependent splicing regulation by TDP-43. Nat Neurosci 14, 452–458 (2011).

27 Wang, W. et al. The inhibition of TDP-43 mitochondrial localization blocks its neuronal toxicity. Nat Med 22, 869–878 (2016).

28 Kuo, P. H., Doudeva, L. G., Wang, Y. T., Shen, C. K. & Yuan, H. S. Structural insights into TDP-43 in nucleic-acid binding and domain interactions. Nucleic Acids Res 37, 1799–1808 (2009).

29 Romani, A. M. P. in Magnesium in the Central Nervous System (eds R. Vink & M. Nechifor) (2011).

30 Smola, M. J., Rice, G. M., Busan, S., Siegfried, N. A. & Weeks, K. M. Selective 2’-hydroxyl acylation analyzed by primer extension and mutational profiling (SHAPE-MaP) for direct, versatile and accurate RNA structure analysis. Nat Protoc 10, 1643–1669 (2015).

31 Guo, S. K., Nan, F., Liu, C. X., Yang, L. & Chen, L. L. Mapping circular RNA structures in living cells by SHAPE-MaP. Methods 196, 47–55 (2021).

32 Danaee, P. et al. bpRNA: large-scale automated annotation and analysis of RNA secondary structure. Nucleic Acids Res 46, 5381–5394 (2018).

33 Mann, J. R. et al. RNA Binding Antagonizes Neurotoxic Phase Transitions of TDP-43. Neuron 102, 321–338 e328 (2019).

34 Kuo, P. H., Chiang, C. H., Wang, Y. T., Doudeva, L. G. & Yuan, H. S. The crystal structure of TDP-43 RRM1-DNA complex reveals the specific recognition for UG- and TG-rich nucleic acids. Nucleic Acids Res 42, 4712–4722 (2014).

35 Moreno, F. et al. A novel mutation P112H in the TARDBP gene associated with frontotemporal lobar degeneration without motor neuron disease and abundant neuritic amyloid plaques. Acta Neuropathol Commun 3, 19 (2015).

36 Garcia Morato, J., et al. Sirtuin-1 sensitive lysine-136 acetylation drives phase separation and pathological aggregation of TDP-43. Nat Commun 13, 1223 (2022).

37 Cohen, T. J. et al. An acetylation switch controls TDP-43 function and aggregation propensity. Nat Commun 6, 5845 (2015).

38 Cao, Q., Boyer, D. R., Sawaya, M. R., Ge, P. & Eisenberg, D. S. Cryo-EM structures of four polymorphic TDP-43 amyloid cores. Nat Struct Mol Biol 26, 619–627 (2019).

39 Wang, A. et al. A single N-terminal phosphomimic disrupts TDP-43 polymerization, phase separation, and RNA splicing. EMBO J 37 (2018).

40 Oiwa, K. et al. Monomerization of TDP-43 is a key determinant for inducing TDP-43 pathology in amyotrophic lateral sclerosis. Sci Adv 9, eadf6895 (2023).

41 Saibil, H. Chaperone machines for protein folding, unfolding and disaggregation. Nat Rev Mol Cell Biol 14, 630–642 (2013).

42 Yang, X. W. et al. MutS functions as a clamp loader by positioning MutL on the DNA during mismatch repair. Nat Commun 13, 5808 (2022).

43 Gasset-Rosa, F. et al. Cytoplasmic TDP-43 De-mixing Independent of Stress Granules Drives Inhibition of Nuclear Import, Loss of Nuclear TDP-43, and Cell Death. Neuron 102, 339–357 e337 (2019).

44 Lu, S. et al. Heat-shock chaperone HSPB1 regulates cytoplasmic TDP-43 phase separation and liquid-to-gel transition. Nat Cell Biol 24, 1378–1393 (2022).

45 Schlich, M., et al. Cytosolic delivery of nucleic acids: The case of ionizable lipid nanoparticles. Bioeng Transl Med 6, e10213 (2021).

46 Joseph, B. J. et al. TDP-43-dependent mis-splicing of KCNQ2 triggers intrinsic neuronal hyperexcitability in ALS/FTD. Nat Neurosci 28, 2476–2492 (2025).

47 Seddighi, S. et al. Mis-spliced transcripts generate de novo proteins in TDP-43-related ALS/FTD. Sci Transl Med 16, eadg7162 (2024).

48 Walker, A. K. et al. Functional recovery in new mouse models of ALS/FTLD after clearance of pathological cytoplasmic TDP-43. Acta Neuropathol 130, 643–660 (2015).

49 Cao, S. M., et al. Chimeric archaeal BHB-human pre-tRNA scaffold HARBOR system enables versatile production of functional circular RNA modalities. BioRxiv (2026).

50 Lacoste, J. et al. Pervasive mislocalization of pathogenic coding variants underlying human disorders. Cell 187, 6725–6741 e6713 (2024).

51 Ittner, L. M. & Gotz, J. Amyloid-beta and tau--a toxic pas de deux in Alzheimer’s disease. Nat Rev Neurosci 12, 65–72 (2011).

52 Mehra, S., Sahay, S. & Maji, S. K. alpha-Synuclein misfolding and aggregation: Implications in Parkinson’s disease pathogenesis. Biochim Biophys Acta Proteins Proteom 1867, 890–908 (2019).

53 Wu, M., Yang, L. Z. & Chen, L. L. Long noncoding RNA and protein abundance in lncRNPs. RNA 27, 1427–1440 (2021).

54 Elguindy, M. M. & Mendell, J. T. NORAD-induced Pumilio phase separation is required for genome stability. Nature 595, 303–308 (2021).

55 Li, Z. et al. Allele-selective lowering of mutant HTT protein by HTT-LC3 linker compounds. Nature 575, 203–209 (2019).

56 Teng, F. et al. Efficient amyloid-beta degradation in Alzheimer’s disease using SPYTACs. Cell (2026).

57 Xie, M. et al. TREM2 interacts with TDP-43 and mediates microglial neuroprotection against TDP-43-related neurodegeneration. Nat Neurosci 25, 26–38 (2022).

58 Delaglio, F. et al. NMRPipe: a multidimensional spectral processing system based on UNIX pipes. J Biomol NMR 6, 277–293 (1995).

59 Lee, W., Tonelli, M. & Markley, J. L. NMRFAM-SPARKY: enhanced software for biomolecular NMR spectroscopy. Bioinformatics 31, 1325–1327 (2015).

60 Senavirathne, G. et al. Widespread nuclease contamination in commonly used oxygen-scavenging systems. Nat Methods 12, 901–902 (2015).

61 Yuan, J., He, K., Cheng, M., Yu, J. & Fang, X. Analysis of the steps in single-molecule photobleaching traces by using the hidden markov model and maximum-likelihood clustering. Chem Asian J 9, 2303–2308 (2014).

62 Liu, Y. X., Jiang, B. W., Liu, K. M., Yang, L. & Chen, L. L. abcFISH enables multiplexed, single-molecule visualization of circular RNA spatial heterogeneity. BioRxiv (2025).

63 Bolger, A. M., Lohse, M. & Usadel, B. Trimmomatic: a flexible trimmer for Illumina sequence data. Bioinformatics 30, 2114–2120 (2014).

64 Langmead, B., Trapnell, C., Pop, M. & Salzberg, S. L. Ultrafast and memory-efficient alignment of short DNA sequences to the human genome. Genome Biol 10, R25 (2009).

65 Kim, D., Paggi, J. M., Park, C., Bennett, C. & Salzberg, S. L. Graph-based genome alignment and genotyping with HISAT2 and HISAT-genotype. Nat Biotechnol 37, 907–915 (2019).

66 Liao, Y., Smyth, G. K. & Shi, W. featureCounts: an efficient general purpose program for assigning sequence reads to genomic features. Bioinformatics 30, 923–930 (2014).

67 Ritchie, M. E. et al. limma powers differential expression analyses for RNA-sequencing and microarray studies. Nucleic Acids Res 43, e47 (2015).

68 Wang, Y. et al. rMATS-turbo: an efficient and flexible computational tool for alternative splicing analysis of large-scale RNA-seq data. Nat Protoc 19, 1083–1104 (2024).

